# Characterization of norbelladine synthase and noroxomaritidine/norcraugsodine reductase reveals a novel catalytic route for the biosynthesis of Amaryllidaceae alkaloids including the Alzheimer’s drug galanthamine

**DOI:** 10.1101/2022.07.30.502154

**Authors:** Bharat Bhusan Majhi, Sarah-Eve Gélinas, Natacha Mérindol, Isabel Desgagné-Penix

## Abstract

Amaryllidaceae alkaloids (AAs) are a large group of pharmacological plant-specialized metabolites. Norbelladine is the entry compound in AAs biosynthesis. Two enzymes are capable of catalyzing this reaction *in-vitro*, both with low-yield; 1) norbelladine synthase (NBS) condenses tyramine and 3,4-dihydroxybenzaldehyde, while 2) noroxomaritidine/norcraugsodine reductase (NR), catalyzes a reduction reaction. To clarify the mechanisms involved in this controversial step, both *NBS* and *NR* were characterized from *Narcissus papyraceus* and *Leucojum aestivum*. Assays with each enzyme suggested that NBS and NR function together for norbelladine formation. Using molecular-homology modeling and docking studies, we predicted models for the binding of substrates to NBS and NR. Moreover, NBS and NR physically interact, localize to the cell cytoplasm and nucleus and are expressed at high levels in bulbs. Our study establishes that both NBS and NR participate in the biosynthesis of norbelladine, catalyzing the first key steps involved in the biosynthesis of the Alzheimer’s drug galanthamine.

## INTRODUCTION

Amaryllidaceae alkaloids (AAs) are a large group of plant specialized metabolites with large therapeutical potentials. The greatest commercial success among AAs is galanthamine, produced by many *Narcissus, Galanthus* and *Leucojum* species, and currently used as an acetylcholinesterase inhibitor to fight Alzheimer’s disease symptoms^1^. Other AAs with strong antiviral activity, such as lycorine and cherylline, are intensively studied to fight emerging infectious diseases^2–4^. AAs have complex carbon skeleton and are challenging to chemically synthesize. Hence, they are often extracted directly from plants, limiting their broad usage due to the often low and variable quantity produced *in vivo*. Their massive extraction would possibly mean a loss in the biodiversity of the endogenous flora of some countries. One interesting alternative would be to biosynthesize them in host microorganisms, developing a sustainable platform of production, but this requires prior knowledge of the metabolic pathway. Ironically, much more is known about the pharmacology of AAs than about their biosynthesis. Few genes encoding biosynthetic enzymes are known and scarcely any of the known enzymatic reactions have been fully characterized^5^.

Despite their diverse chemical structures, AAs share norbelladine as a common biosynthetic origin (Figure 1). Following its synthesis, norbelladine undergoes several chemical modifications achieved by a multitude of enzymes catalyzing various types of reactions, such as *O-* and *N-*methylations (OMTs, NMTs), C-C and C-O bond formations, oxidations and reductions, demethylations, and hydroxylations resulting in a variety of different structural types of AAs^6–11^.

**Figure 1.**
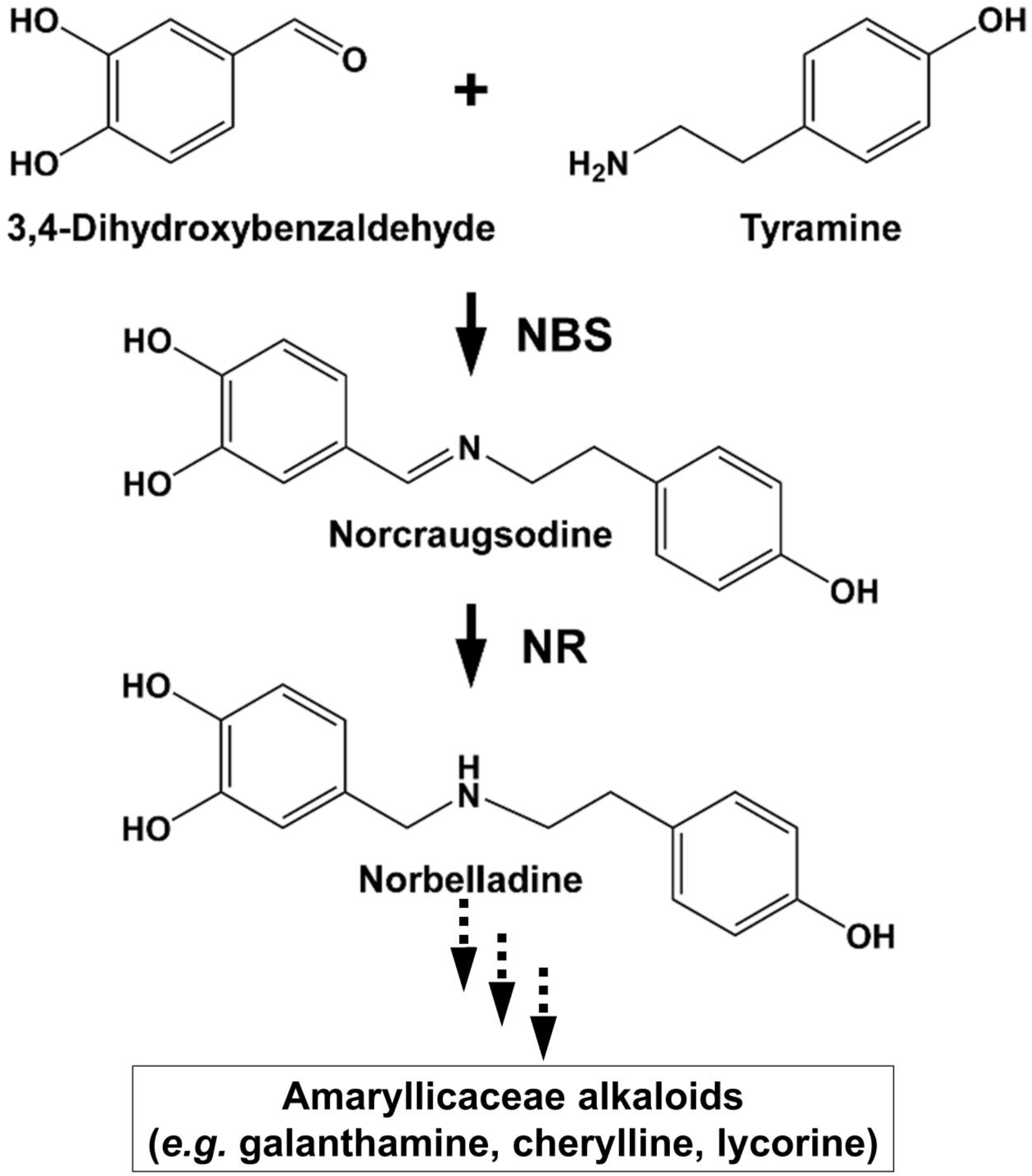
Norbelladine synthase (NBS) and noroxomaritidine/norcraugsodine reductase (NR) is involved in the proposed biosynthetic pathway for galanthamine, lycorine and haemanthamine. NBS catalyzes the condensation of tyramine and 3,4-dihydroxybenzaldehyde (3,4-DHBA) to form norcraugsodine and noroxomaritidine/norcraugsodine reductase (NR) reduces it to form norbelladine, the common precursor to all Amaryllidaceae alkaloids produced in plants including galanthamine, lycorine and haemanthamine.

Previous studies have tried to unravel norbelladine biosynthesis because of its pivotal role in AAs biosynthesis. We and others identified several AA biosynthetic genes encoding enzymes involved in these early steps^11–16^. Norbelladine originates from the condensation of tyramine and 3,4-dihydroxybenzaldehyde (3,4-DHBA), both derived from the amino acids L-tyrosine and L-phenylalanine, respectively. The enzymes responsible for tyramine and 3,4-DHBA biosynthesis are known and spread in most plant species (*e.g.,* tyrosine decarboxylase (TYDC), phenylalanine ammonia-lyase (PAL), cinnamate 4-hydroxylase (C4H), *etc.*). The enzymes and intermediates implicated in norbelladine synthesis are less clear and specific to Amaryllidaceae. The currently accepted model states that the condensation of tyramine and 3,4-DHBA leads to the formation of the imine norcraugsodine, further reduced to produce norbelladine^17^. To date, there are two reported enzymes capable of catalyzing these reactions *in vitro*, but both perform it with very low yield^12,14^. The first one is norbelladine synthase (NBS), recently characterized from two plant species *N. pseudonarcissus* (*NpKA*NBS) and *L. aestivum* (*La*NBS). NBS was shown to condense tyramine and 3,4-DHBA to form low levels of norbelladine^14,16^. The second one, noroxomaritidine/norcraugsodine reductase (NR) from *N. pseudonarcissus* (*NpKA*NR) mainly catalyzes noroxomaritidine synthesis, but also produces above background levels of norbelladine following incubation with tyramine, 3,4-DHBA and NADPH^12^. Therefore, the contribution of these two enzymes to norbelladine synthesis *in vivo* is still not clear.

We hypothesized that norbelladine is formed through two separate reactions (condensation and reduction) catalyzed by two different enzymes (Figure 1). In this study, we characterized the NBS and NR enzymes from two Amaryllidaceae plants species *N. papyraceus* and *L. aestivum* to elucidate the crucial reactions leading to norbelladine synthesis. We propose that for NBS or NR to synthesize norbelladine efficiently, this step optimally requires a two-step reaction catalyzed by both enzymes interacting together in a metabolon.

## RESULTS

### Identification and structure analysis of NBS from *N. papyraceus* and *L. aestivum*

The full-length cDNAs of *NpNBS*, *LaNBS*, *NpNR*, and *LaNR* genes were obtained from previously reported transcriptome sequencing of *N. papyraceus* and *L. aestivum*^15,16^. The open reading frame (ORF) of both *NpNBS* and *LaNBS* gene is 480 bp and encodes a 159-amino acid protein (Figure S1). *In silico* protein analysis by DNAMAN analysis software indicated that *Np*NBS and *La*NBS had a predicted molecular weight (MW) of 17.4 kDa and a theoretical isoelectric point (pI) of 5.3 and 5.1, respectively. Multiple amino acid sequence alignments of *Np*NBS and *La*NBS with the already characterized protein *NpKA*NBS reaction^14^ and norcoclaurine synthase from *T. flavum* (*Tf*NCS)^18^ showed that *Np*NBS and *La*NBS share over 41% of amino acids sequence identity with the ortholog *Tf*NCS (Figure S1). In addition, *NpKA*NBS, *Np*NBS and *La*NBS share 83% identity between each other, while *Np*NBS and *La*NBS share 85% of identity (Figure S1). Domain search using *NCBI-conserved-domain-search service* revealed the presence of conserved Bet v1 and Pathogenesis-Related (PR-10) protein domains in both *Np* and *La*NBS. Both contained the phosphate-binding loop (P-loop) glycine-rich region (Figure S1), a conserved ligand-binding domain of Bet v1 protein family^19^. As reported previously, the alignment showed that *Tf*NCS catalytic residues Tyr108, Glu110 and Lys122 are well-conserved in *Np* and *La*NBS, corresponding to Tyr68, Glu71, and Lys83, respectively, in their sequences (Figure S1)^14,18^. During norcoclaurine synthesis, *Tf*NCS Glu110 and Lys122, two strong proton exchangers, directly catalyze the reaction, and Tyr108, as a hydrogen-binding donor, rather contributes to the electrostatic properties of the active site and defines the shape of the cavity entrance^18^. The fourth catalytic residue Asp141 from *Tf*NCS is replaced by hydrophobic Ile102 in both *Np* and *La*NBS, as reported previously for *NpKA*NBS^14^. Homology modeling of the enzymes from both *Np* and *La* revealed a striking structure similarity with superimposed template crystal of *Tf*NCS, analogous to the overall structure of Bet v1-like proteins family (Figure 2a, Figure S2a, and Figure S3a-d). As *Tf*NCS, both NBS are composed by seven-stranded antiparallel β-sheets, two long C- and N-terminal helices and two short ones, enclosing a cleft with polar residues at its entrance and hydrophobic residues in its core (Figure 2a, and Figure S2a). Using MOE’ Site Finder tool, *Np* and *La* NBS cavity is predicted to contain an active site of 27 and 37 residues, respectively (Table S1). For both *Np* and *La*NBS, the cavity is surrounded by the catalytic residues Tyr68, Glu71, Lys83 and Ile102, with Tyr68 at its entrance, Lys83 at the other side and the P-loop at the bottom (Figure 2b, and Figure S2b).

**Figure 2.**
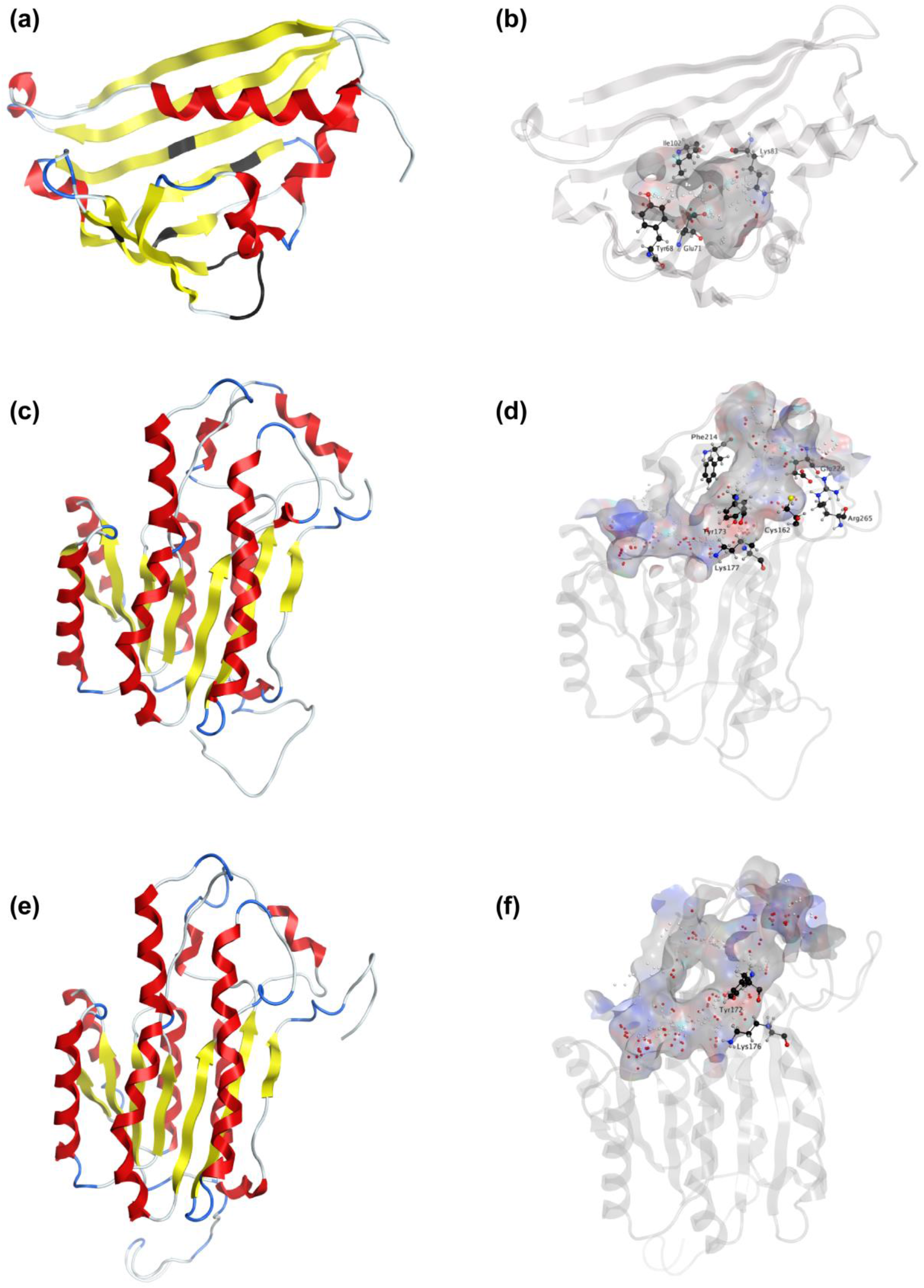
Homology modeling of *Np*NBS, NR and TR. (a) Ribbon representation of *Np*NBS with colored secondary structures. β-sheets are displayed yellow, α-helices red, and loops blue. Conserved glycine rich P-loop of PR10-/Betv1 enzymes is shown black, as are conserved active site residues of norcoclaurine synthase Tyr68 (Tyr108 from *Tf*NCS), Glu71 (Glu111), Lys83 (Lys123). (b) Transparent ribbon representation of *Np*NBS with predicted active site pocket in transparent surface. The predicted ligands site computed by the Site Finder tool of MOE software is displayed as white and red alpha sphere centers inside the pocket. Catalytic residues Tyr68, Glu71, Lys83 as well as residue Ile102 all surrounding the active site are shown as black sticks. (c) Ribbon representation of *Np*NR with highlighted secondary structures. β-sheets are displayed yellow, α-helices red and loops in blue. (d) Cartoon ribbon representation with transparent surface view of the predicted active site forming a catalytic tunnel that crosses the enzyme. The predicted ligands site is displayed as white and red alpha sphere centers inside the pocket. Conserved active site residues Cys162, Tyr173, Lys177, Phe214, Glu224, Arg265 surrounding the tunnel are shown as black sticks. (e) Ribbon representation of *Np*TR with secondary structures. β-strands are displayed yellow, α-helices red, and loops blue. (f) Ribbon representation with transparent surface view of predicted active site tunnel crossing the enzyme. Predicted ligands site is displayed as white and red alpha sphere centers. Conserved catalytic Tyr172 and Lys176 are shown as black sticks.

### Identification and structure analysis of NR from *N. papyraceus* and *L. aestivum*

The ORF of the *NpNR* and *LaNR* genes is 810 bp, both encoding a 269-amino acid protein (Figure S4). *Np*NR and *La*NR have a predicted molecular weight (MW) of 29 kDa and theoretical pI of 6.0 and 5.4, respectively. Sequence comparison and domain search confirmed the presence of conserved short-chain dehydrogenases/reductases (SDR) domain in *Np* and *La*NR. *NpKA*NR, *Np*NR and *La*NR share over 76% of identity with each other, while *Np*NR and *La*NR share 90% of identity (Figure S4). Like all classical SDRs, *Np*NR and *La*NR contain a TGXXX[AG]XG cofactor binding motif and a YXXXK active site motif, with the Tyr and Lys of the active site serving as critical catalytic residues (Figure S4). Structurally, NRs predicted models from both species are formed by a seven-stranded parallel b-sheet inserted between a pair of three a-helices (Figure 2c, and Figure S2c). A long tunnel is shaped at the C-termini of b-strands partially wrapped by the a-helices and loops that elevate beyond the b-sheets (Figure 2c, and Figure S2c). *Np* and *La*NR tunnel active site are predicted to contain 55 and 63 residues, respectively (Table S1). A strong similarity in structure of both *Np* and *La*NR with *N. pseudonarcissus* noroxomaritidine/norcraugsodine reductase (*NpKA*NR, 5FF9) was noted (Figure S5a-d). In general, amino acids involved in NADPH binding by noroxomaritidine/norcraugsodine reductase (*NpKA*NR, 5FF9) are conserved and similarly oriented, *i.e.* Val81 (Val83 for *NpKA*NR); Asp80 (82), Arg55 (57), Ser54 (56), Thr32 (34), catalytic residue Tyr173 (175), catalytic residue Lys177 (179), Asn108 (110); Thr208 (210), Pro203 (205), Gly204 (206) and Ala205 (207). At the site of ligand interaction, the tunnel expands into a larger pocket where aromatic Phe214 (216) is conserved, Ala112 replaces Tyr114, both possibly involved in polycyclic substrate orientation and binding. At the extremity, Glu224 (226), Arg265 (267), Cys162 (164), His170 (172) are preserved. This overall similarity is also reflected by the striking superimposition of the two proteins secondary and tertiary structures with crystalized *NpKA*NR (Figure S5b and d).

To identify other reductases that could catalyze similar reduction reaction, we searched for homologs of NR in the transcriptome sequences of *N. papyraceus*^15^ and *L. aestivum*^16^, and identified a tropinone reductase (TR) belonging to the same SDR superfamily in both species. The ORF of the *NpTR* is 825 bp and encodes a 274-amino acid protein while the *LaTR* is 816 bp and encodes a 271-amino acid protein (Figure S4). *Np*TR and *La*TR have a predicted molecular weight (MW) of 30 kDa and 29.5 kDa and theoretical pI of 6.9 and 8.4, respectively. *Np*TR and *La*TR contain the TGXXX[AG]XG cofactor binding motif and the YXXXK catalytic active site motif (Figure S4). Gly206 is replaced by a tryptophan at position 203 and 200 in *Np* and *La*TR, respectively, although this residue was conserved in active SDR/tropinone reductases^20^. Multiple sequence alignments showed that *NpKA*NR, *Np*NR, *La*NR, *Np*TR, and *La*TR share over 58% of identity between each other. Predicted TRs structures are similar to NRs with some key differences in the ligand active site, including replacement of Phe216 from *NpKA*NR by Arg213 in *Np*TR and by Leu210 in *La*TR, while Tyr114 is replaced by Asn111 in *Np*TR and Asn108 in *La*TR (Figure 2e,f, Figure S2e,f, S6, and Table S1).

### Predicted interactions of ligands with NBS and NR

To shed light on the reactions involved in norbelladine synthesis (Figure 1), we studied the interaction of NBS and NR with their respective proposed ligands through molecular docking analysis *in silico*. Tyramine and 3,4-DHBA were docked with scores -5.03 and -5.23 kCal/mol inside the NBS pocket (Figure 3, Figure S7, and Table S1). Most of the obtained poses displayed the same ligands orientation where 3,4-DHBA and tyramine adopted a stack configuration with respective aromatic rings lying on near-to-parallel planes, similarly to dopamine and hydroxybenzaldehyde in crystalized *Tf*NCS (2VQ5) (Figure 3a,b, Figure S7a,b). The carbonyl group of 3,4-DHBA faced the amine group of tyramine. Lys83, whose proposed role is to intercept the carbonyl group of the aldehyde substrate, interacted with 3,4-DHBA carbonyl end through a hydrogen bond (Figure 3b, and Table S1). At the other end, the hydroxyl group of C4 was hydrogen bonded with the possibly base-acting residue Glu71. All predicted poses implied interaction with Glu71 and Lys83 strengthening the probability of their key role in the catalytic mechanism. Tyramine was hold in place by stacking interaction, and a hydrogen bond between its amine and 3,4-DHBA carbonyl group. PLIP software^21^ predicted additional hydrophobic interactions between Phe73, Thr85, Phe104 and Ile143, and 3,4-DHBA, as well as two hydrogen bonds of tyramine with Ser31 and Tyr59 (Figure 3b, Figure S7b, and Table S1). In general, these predicted interactions and spatial arrangements of the ligands inside NBS are consistent with the reaction proposed by Ilari *et al*. in 2009 that would lead to norcraugsodine biosynthesis.

**Figure 3.**
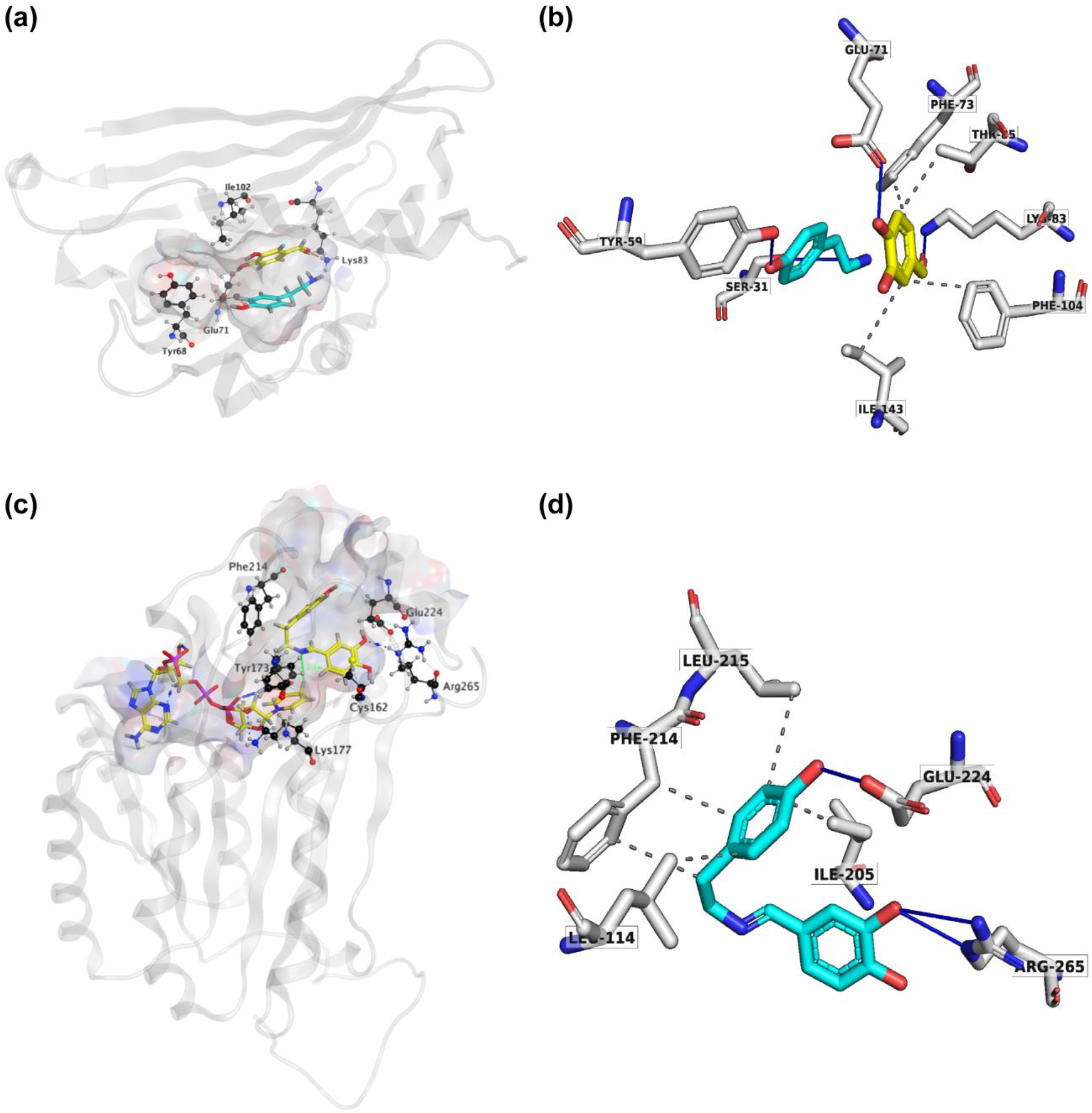
*Np*NBS docked with 3,4-dihydroxybenzaldehyde (3,4-DHBA) and tyramine. (a) Cartoon representation of *Np*NBS with transparent surface-active site pocket docked with 3,4-DHBA (up) and tyramine (down). Conserved catalytic residues Tyr68 (Tyr108 from *Tf*NCS), Glu71 (Glu110), Lys83 (Lys122) and Ile102 (instead of Asp141) shaping the binding site are shown as black sticks. (b) PLIP predicted conformation of interacting residues of *Np*NBS (grey) docked with 3,4-DHBA (turquoise) and tyramine (yellow). (c) *Np*NR docked with NADPH and norcraugsodine. Ligands NADPH (left) and norcraugsodine (right) are represented as thick yellow sticks. Conserved active site residues Cys162, Tyr173, Lys177, Phe214, Glu224 and Arg265) shaping the binding site are shown as thin black sticks. (d) PLIP predicted interacting residues of docked norcraugsodine (turquoise) with *Np*NR (grey sticks).

Following its formation, norcraugsodine would be transferred from NBS to NR active site to be reduced into norbelladine (Figure 1) with NADPH as the electron donor. Upon docking, NADPH positioned into *La* and *Np*NR modeled active site following a similar arrangement compared to crystalized reductases such as 5FF9 (*NpKA*NR) with a score of -11.51 kCal/mol (Figure 3c,d, and Figure S7c,d, and Table S1). It interacted with Gly36, Cys53, Arg55, Val109, Ile35, His170, Thr111, Lys177, Asn108, Thr206. PLIP confirmed these interacting residues and additionally predicted hydrophobic interaction with Ile35, hydrogen-bonds with Lys33, Cys79, Gly110, Thr208, Val209 and Gln210; π-Cation interactions and salt bridges with Arg55 (Table S1). Close to the ligand-binding site, the nicotinamide ring faces Ile158, Pro203, Gly204 on one side, and the substrate-binding pocket on the B-side. As was the case for docked noroxomaritidine in the active site of noroxomaritidine reductase, docking predicted that norcraugsodine binds to the active site of NR by a combination of polar and non-polar interactions. Docked norcraugsodine displayed two possible conformations in *Np* and *La* NR: either bended or diagonal. For both conformations, the amine group of norcraugsodine was positioned close to C4 of NADPH and close to Tyr173, obtaining a docking score of -6.36 and - 6.15 kCal/mol (*NpNR* and *La*NR, respectively, Figure 3c,d, Figure S7c,d, and Table S1). In both cases, norcraugsodine phenol cycle was situated near Phe214, and its dihydroxybenzene group positioned close to Glu224, interacting with His170 and Arg265. PLIP predicted additional hydrophobic interactions of norcraugsodine with Leu114, Ile205, Phe214 and Leu215, and hydrogen-bonding with Glu224 (Figure 3c,d, Figure S7c,d, and Table S1). These interactions are consistent with the proposed reduction of the nitrogen-carbon double bond of norcraugsodine to yield norbelladine involving NADPH and the catalytic residues Tyr173 and Lys177.

### NBS and NR produce higher titers of norbelladine together than separately

To examine the NBS, NR, and TR protein function from both *N. papyraceus* and *L. aestivum*, the full-length ORFs were PCR-amplified from *N. papyraceus* and *L. aestivum* bulb cDNA, cloned and expressed proteins were purified (Figure S8). We were unable to purify the *La*TR enzyme in our experimental conditions. We first tested the NBS, and NR purified proteins from *Np* and *La* separately in assays containing tyramine, 3,4-DHBA, and NADPH. The resulting assay products were subjected to HPLC-MS/MS analysis using Positive-Electrospray-Ionization-mode (ESI+). Before injecting the assays, norbelladine standard was injected at 1 mg/L and predicted major mass spectral parent-ion for the norbelladine m/z 260 [M + H]+ at 3.435 min (Figure 4a, and Figure S9a-r) was observed. Fragmentation of the norbelladine parent-ion m/z 260 yielded to major ion fragments of m/z 121, 123 and 138 using 10 V as collision energy. Multiple-reaction-monitoring (MRM) transitions of 260 → 138 m/z and 260 → 121 m/z were selected, optimized, and used as quantifier and qualifier ions respectively. We could not observe any parent-ion mass for the norcraugsodine standard, despite repeated trials. We inferred that norcraugsodine is highly unstable in solution and/or thermolabile so the heat used during the ionization process in the HPLC-MS/MS source could lead to its degradation. Enzyme assays containing recombinant NBS protein from *Np* or *La*, 3,4-DHBA, tyramine, and without or with NADPH yielded a peak at 3.435 min in MRM-acquisition mode on HPLC-MS/MS which was the same retention time and MRM transitions as those of the norbelladine standard (Figure 4a, and Figure S9e,f). Similarly, for the assays examining the NR, tyramine and 3,4-DHBA were incubated with NR and NADPH and the resulting product showed a peak with the same retention time as norbelladine standard in the reaction mixture of *Np* enzyme or *La* enzyme (Figure 4a, and Figure S9g,p). Comparison of the mass spectrum obtained after the fragmentation of authentic norbelladine standard using collision-induced-dissociation (CID) with 10V and the one of the product obtained from NBS and NR reactions showed the fragmentation pattern is the same for both confirming the identity of the enzymatic product (Figure S10). In assays lacking substrates or enzyme, no norbelladine formation was detected (Figure 4a, and Figure S9b,d). Similarly, MBP tag alone protein purified from *E. coli* transformed with empty pMAL-c2X vector showed no activity (Figure 4a, and Figure S9c). As expected, NR assays lacking NADPH resulted in no norbelladine production (Figure 4a, and Figure S9c). As reported before^12,14,16^, our results confirm that both NBS and NR alone from *N. papyraceus* and *L. aestivum* is able to catalyze the reaction for condensation/reduction of tyramine and 3,4-DHBA to produce a lower amount of norbelladine (Figure 4a).

**Figure 4.**
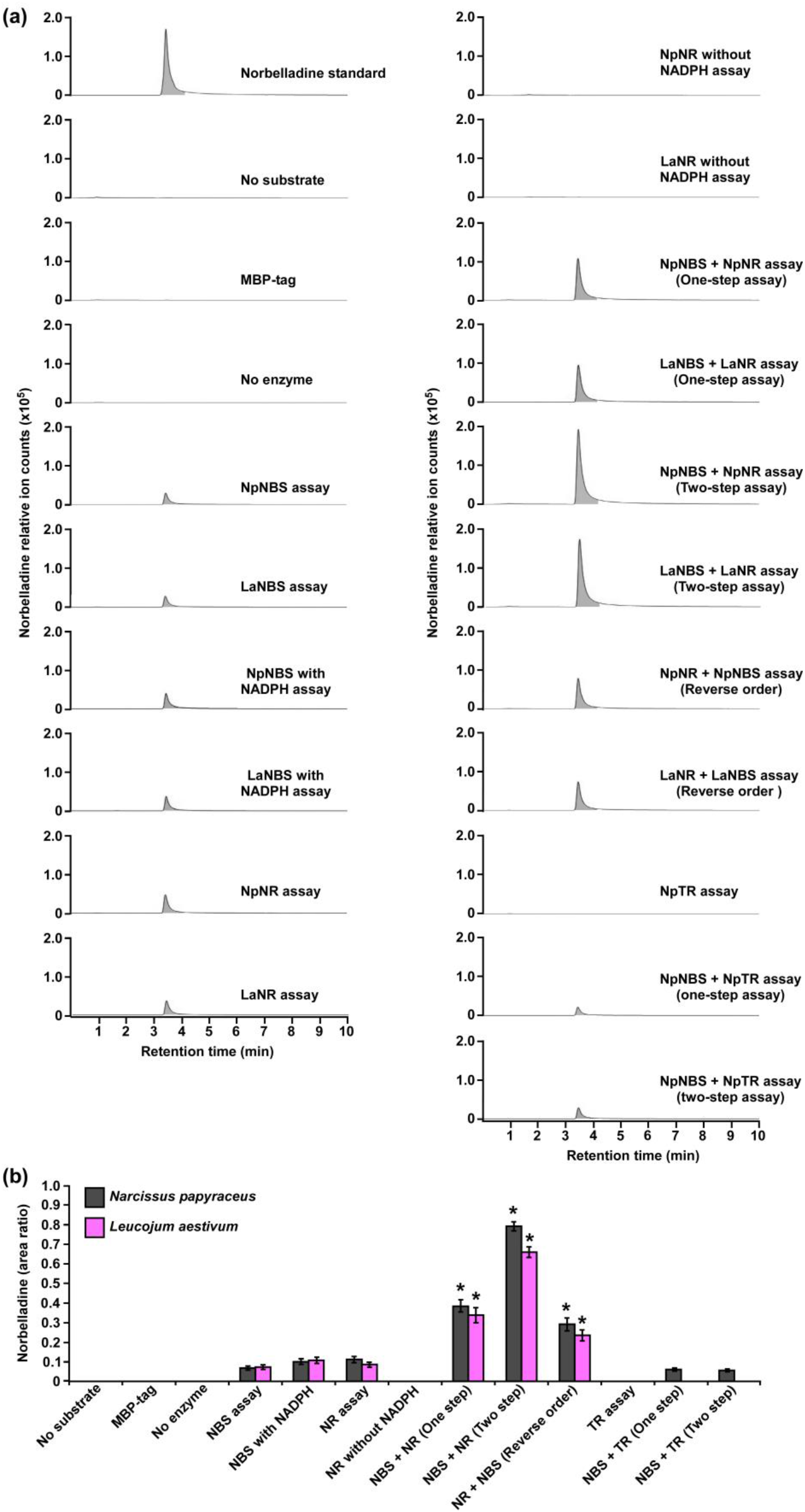
Enzymatic activity of NBS, NR and TR. Enzymes were tested separately or together for production of norbelladine, and the reaction product was monitored using HPLC-MS/MS. (a) Extracted ion chromatograms of quantifier MRM transition 260 → 138 m/z showing the product norbelladine in different enzymatic assays. The tested substrates used were 3,4-DHBA (300 μM) and tyramine (10 μM), and panels show norbelladine standard; assay without substrates; assay with MBP tag; assay without enzyme; and the complete assay performed with recombinant *Np*NBS, *La*NBS, *Np*NR, *La*NR and NpTR recombinant enzymes as indicated. Parent ion mass-to-charge (m/z) of 260 for norbelladine was subjected to collision-induced dissociation using multiple reaction monitoring (MRM) analysis. (b) Comparison and relative quantification of assays shown in Figure 4a in triplicate (mean ± SD, n = 3). The norbelladine product profiles in different assays performed were analyzed by HPLC-MS/MS and the obtained amount were quantified using the area ratio of norbelladine produced in the assay to the papaverine internal standard. Data are means ± *SE* of three biological repeats. Asterisks indicate a significant difference (Student’s *t* test, *p* < 0.05) relative to *Np*NBS alone enzymatic assay.

To examine if both NBS and NR work together for the condensation of tyramine and 3,4-DHBA into norcraugsodine followed by a reduction into norbelladine (Figure 1), we tested NBS, and NR purified enzymes together in one-step assay or in two-step sequential manner. Our inability to detect the intermediate norcraugsodine through HPLC-MS/MS prompted us to examine the norbelladine production in each assay mixture. We measured the substrates and equal amount of added papaverine (internal standard) in all reaction mixture for accurate quantification of the observed product (Figure S9a-r). Norbelladine production was significantly (six-fold) higher when both *Np*NBS and *Np*NR were present in a single reaction, compared to assays with *Np*NBS or *Np*NR separately (Figure 4a,b and Figure S9e-j). Similarly, we observed four-fold higher norbelladine production when *La*NBS and *La*NR were present in a single reaction, compared to assays with *La*NBS or *La*NR individually (Figure 4a,b, and Figure S9o-r). When the reactions were performed in a two-step manner: NBS first followed by NR, we observed two-fold higher norbelladine level than when enzymes were together in a single-step for both *Np* and *La* enzymes (Figure 4a,b and Figure S9i,j,q,r), which was 12 fold (*Np*) and 8 fold (*La*) higher in comparison to assays with the enzymes individually (Figure 4a,b). In the reverse order stepwise reaction for both species, NR first followed by NBS yielded lower levels of norbelladine than observed in the reaction order NBS first followed by NR, but still producing higher amounts compared to single enzyme reactions (Figure 4a,b and Figure S9j,k). Our results indicate that both NBS and NR functions together optimally in a sequential manner (NBS first followed by NR) to produce norbelladine. To further confirm the specificity of NR enzyme in norbelladine production, we tested the *Np*TR purified protein individually and together with *Np*NBS in assays containing 3,4-DHBA, tyramine, and NADPH. We found no norbelladine formation in assays containing *Np*TR enzyme, and assays with both *Np*NBS and *Np*TR in one-step or two-step yielded similar level of norbelladine compared with assays containing only *Np*NBS enzyme (Figure 4a,b and Figure S9l-n). These results confirm the role of NBS and NR together to channel the substrates rapidly and effectively for their condensation and reduction into norbelladine.

### NBS and NR forms dimer and NBS physically interacts with NR *in planta* and in yeast

Previous studies suggest that the norcoclaurine synthase (NCS) proteins appear to assemble as dimers for their catalytic activities^22–24^. Similarly, the NR protein of *N. pseudonarcissus* was shown to exist as a tetramer through crystal structure of the enzyme^12^. Based on these observations, we hypothesized that NBS and NR form dimer for their activity. To explore this possibility, we examined the physical interactions between *Np*NBS-*Np*NBS, *La*NBS-*La*NBS, *Np*NR-*Np*NR, and *La*NR-*La*NR in *N. benthamiana* leaves by split-luciferase-complementation-assays (SLCA). *Np*NBS, *La*NBS, *Np*NR, and *La*NR were fused to the N-terminal half of the luciferase protein (NLuc) and co-expressed via *Agrobacterium* in *N. benthamiana* leaves along with *Np*NBS, *La*NBS, *Np*NR, and *La*NR fused to the C-terminal half of luciferase (CLuc) protein, respectively. As negative controls, *Np*NBS-NLuc, *La*NBS-NLuc, *Np*NR-NLuc, and *La*NR-NLuc were co-expressed with CLuc empty vector and CLuc-*Np*NBS, CLuc-*La*NBS, CLuc-*Np*NR, and CLuc-*La*NR were co-expressed with NLuc empty vector (Figure 5a-d). Expression of all the fusion proteins was validated by western blot analysis (Figure S11a,b). The homodimeric interactions were monitored by measuring luminescence at 48 h after agroinfiltration of the tested protein pairs. Co-expression of *Np*NBS-NLuc with CLuc-*Np*NBS, *La*NBS-NLuc with CLuc-*La*NBS, *Np*NR-NLuc with CLuc-*Np*NR, and *La*NR-NLuc with CLuc-*La*NR resulted in emission of significantly higher luminescence compared to the negative controls indicating a physical interaction *in planta* between NBS-NBS and NR-NR fusion proteins (Figure 5a-d). One explanation for the production of norbelladine in reactions containing NBS and NR is that NR functions as an enzyme that acts on norcraugsodine produced from tyramine and 3,4-DHBA by NBS. Alternatively, NR may alter the catalytic properties of NBS through allosteric regulation, which allows NBS to itself form norbelladine, or vice versa, or it remains formally possible that NBS and NR are present together in a metabolon, NR playing a regulating role guiding the substrates to the imine intermediate and reduction to norbelladine. To test the importance of physical interactions for norbelladine formation, the interactions of *Np*NBS and *La*NBS with full-length *Np*NR and *La*NR were examined in *N. benthamiana* leaves by split-luciferase-complementation-assays. *Np*NBS and *La*NBS were fused to the N-terminal half of the luciferase protein (NLuc) and co-expressed via *Agrobacterium* in *N. benthamiana* leaves along with *Np*NR and *La*NR fused to the C-terminal half of luciferase (CLuc). As a control, *Np*NBS-NLuc and *La*NBS-NLuc were co-expressed with CLuc-*Np*TR and CLuc-*La*TR, respectively. As negative controls, *Np*NBS-NLuc and *La*NBS-NLuc were co-expressed with CLuc empty vector and CLuc-*Np*NR and CLuc-*La*NR were co-expressed with NLuc empty vector (Figure 5a-d). Expression of the examined fusion proteins was confirmed by western blot analysis (Figure S11a,b). Protein-protein interactions *in planta* were quantified by measurements of luminescence at 48 h after agroinfiltration. In agreement with the enzymatic assays, co-expression of *Np*NBS-NLuc with CLuc-*Np*NR and *La*NBS-NLuc with CLuc-*La*NR resulted in emission of significantly higher luminescence compared to the negative controls and the control protein CLuc-*Np*TR and CLuc-*La*TR (Figure 5a-d), demonstrating a physical interaction *in planta* between NBS and NR fusion proteins.

**Figure 5.**
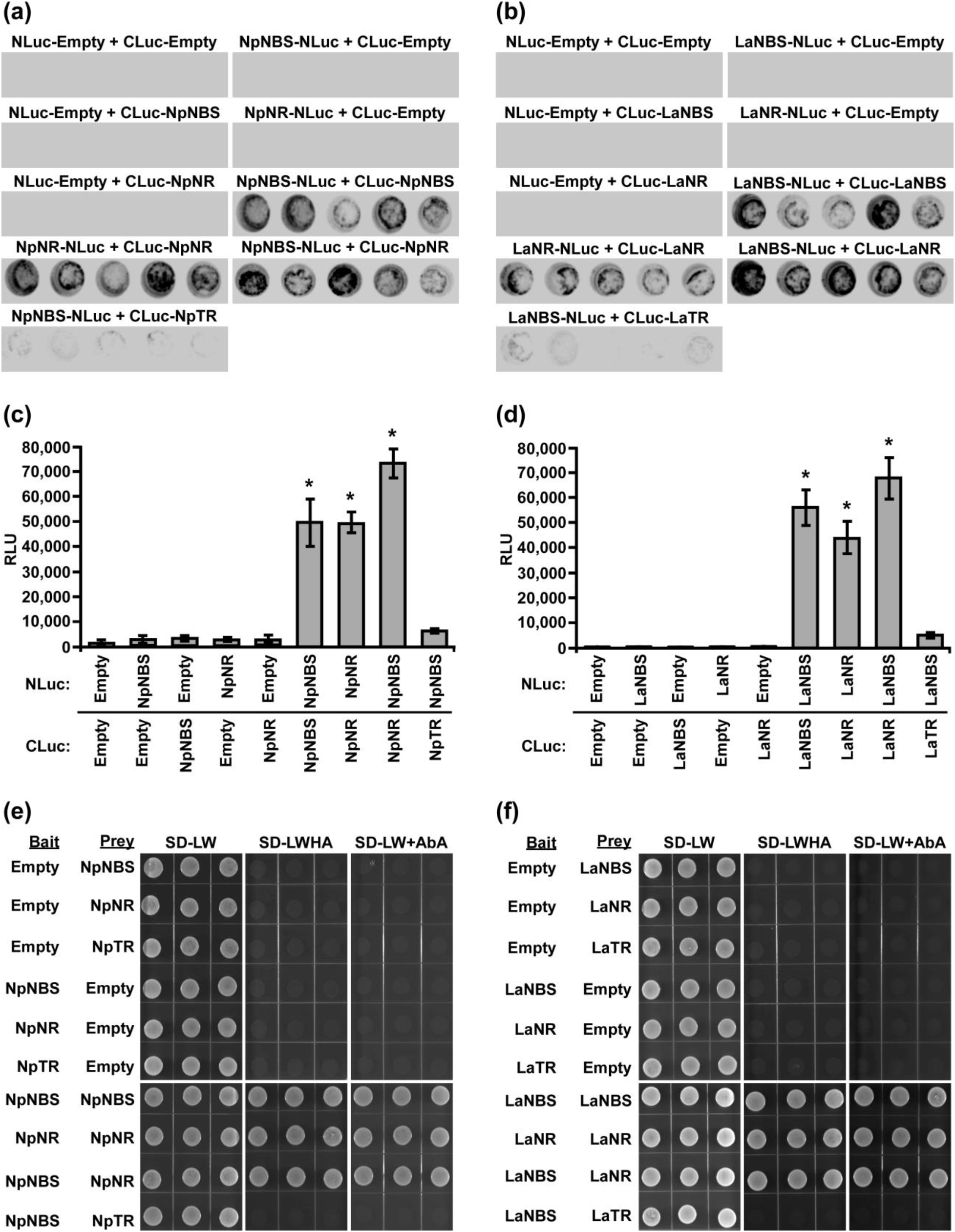
Physical interaction of NBS and NR *in planta* and in Yeast. (a-d) The indicated proteins fused to NLuc or CLuc were expressed in leaves of *Nicotiana benthamiana* plants via *Agrobacterium tumefaciens* infection. (a, b) The images show LUC images of 96 well microtiter plates containing *N. benthamiana* leaf discs expressing the indicated constructs. (c, d) Luciferase activity was quantified as relative luciferase units (RLU) at 48 hr postinfiltration. Data are means ± *SE* of three biological repeats. Asterisks indicate a significant difference (Student’s *t* test, *p* < 0.05) relative to empty vector. (e, f) Yeast expressing the indicated proteins fused to the GAL4 DNA-binding domain (Bait) or to the GAL4 DNA activation domain (Prey) were grown on synthetically defined (SD) medium lacking Leu and Trp (SD-LW), SD-LW lacking His and adenine (SD-LWHA), or SD-LW supplemented with Aureobasidin A (SD-LW+AbA). Empty vectors (EV) were used as negative controls.

To validate the interaction detected *in planta* and to check if the observed interactions are direct, the interaction between NBS-NBS, NR-NR, NBS-NR, and NBS-TR was then examined by yeast two-hybrid system. NBS, NR and TR from both plant species were fused to both bait and prey plasmids. Expression in yeast of bait and prey proteins was confirmed by western blot analysis (Figure S11c,d). Similar interactions between NBS-NBS, NR-NR and NBS-NR were observed (Figure 5e,f) when the same protein pairs from both plant species were expressed in yeast as bait and prey proteins in agreement with the observed interaction *in planta*. As observed *in planta* no interaction was found between NBS-TR in yeast (Figure 5e,f). Taken together, these results obtained in different experimental system indicated the regulatory interactions between NBS and NR and raise the possibility that NBS and NR function as heteromultimeric proteins.

### NBS and NR colocalize in the cell cytoplasm and nucleus

The NBS protein fused with green fluorescent protein (GFP) from both *L. aestivum* and *N. papyraceus* were recently shown to localize to the cell cytoplasm and nucleus^16^. Similarly, we found NR lacked any predicted signal peptides. To investigate NBS and NR subcellular localization, the *NBS* and *NR* coding region from both *N. papyraceus* and *L. aestivum* plants were fused upstream to the yellow-fluorescent-protein gene (*YFP*). The *Np*NBS-YFP, *La*NBS-YFP, *Np*NR-YFP, and *La*NR-YFP fusions were transiently expressed in leaves of *N. benthamiana* plants via *A. tumefaciens*, and their localization was monitored by confocal fluorescence microscopy. The cyan fluorescent protein (CFP), which localizes to the cytoplasm and nucleus^25^, was used as a control. Expression of all the fusion proteins was validated by western blot (Figure S12). As shown in Figure 6a, the NBS-YFP and NR-YFP fusion protein from both plant species localized in the cell cytoplasm and nucleus, like CFP. These results suggest that both NBS and NR are distributed to the same nucleocytoplasm cellular compartment. To further confirm the colocalization pattern of NBS and NR, the NR coding region from both *N. papyraceus* and *L. aestivum* plants were fused upstream to the cyan fluorescent protein (CFP) and co-expressed via *Agrobacterium* in *N. benthamiana* leaves along with *Np*NBS and *La*NBS fused to the YFP, and their localization pattern was monitored by confocal microscopy. Expression of all the fusion proteins was validated by western blot (Figure S12). Similar profiles of fluorescence pattern in the cell cytoplasm and nucleus were observed for NBS-YFP and NR-CFP (Figure 6b). These results confirm that NBS and NR which lack predicted signal peptides, colocalized to both the cytoplasm and nucleus of cell.

**Figure 6.**
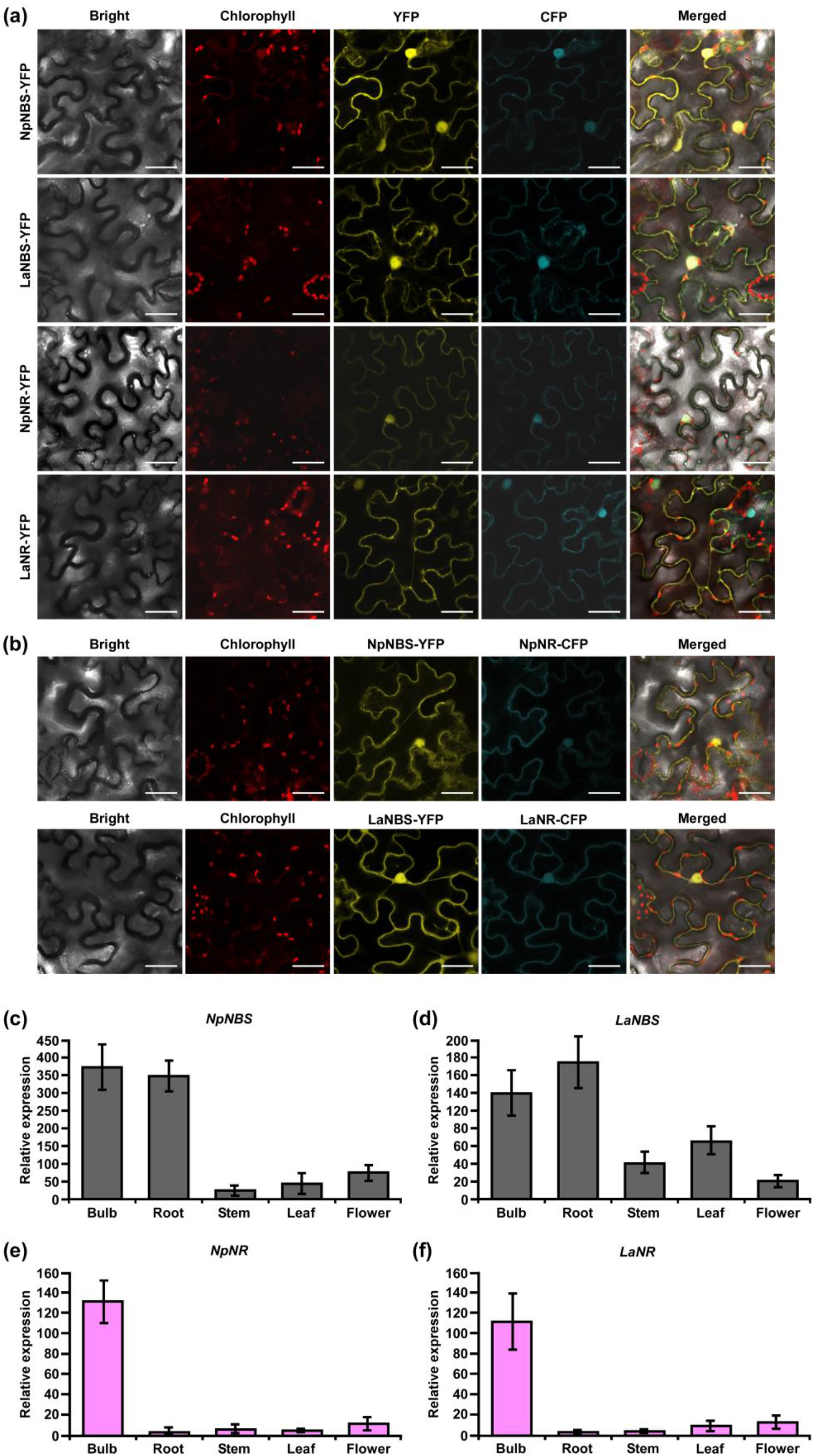
Cellular localization and relative expression of *NBS* and *NR*. (a) The indicated fusion proteins were coexpressed with the cyan fluorescent protein (CFP) in *Nicotiana benthamiana* leaves via *Agrobacterium tumefaciens*. After 48 hr, fluorescence was monitored in epidermal cells by confocal microscopy. Bright field, chlorophyll, yellow fluorescent protein (YFP), cyan fluorescent protein (CFP), and merged fluorescence images are shown. Scale bars in images represent 50 μM. (b) Confocal micrographs of transiently expressed NBS-YFP and NR-CFP in *Nicotiana benthamiana* leaves showing they colocalize to the cytoplasm and nucleus. Bright field, chlorophyll, yellow fluorescent protein (YFP), cyan fluorescent protein (CFP), and merged fluorescence images are shown. Scale bars in images represent 50 μM. (c-f) Relative expression of *NBS*, and *NR* in different tissues of *N. papyraceus* and *L. aestivum* using reverse transcription quantitative PCR (RT-qPCR) analysis. Different plant tissues as indicated were harvested after flowering, and *NpNBS* (c), *LaNBS* (d), *NpNR* (e), *LaNR* (f) mRNA levels were measured by RT-qPCR analysis relative to expression in leaves. *NpHISTONE* and *LaGAPDH* were used as normalizer. Data are means ± *SE* of three biological repeats.

### *NBS* and *NR* are expressed at high levels in the bulbs of *N. papyraceus* and *L. aestivum*

The expression profiles of the *NBS* and *NR* in different tissues including bulb, root, stem, leaf, and flower were evaluated by using quantitative real-time PCR (qRT-PCR) analysis. The results showed that *NBS* was ubiquitously expressed in all tissues detected, with the highest expression levels in bulb and root (Figure 6c,d). The expression patterns of *NpNBS* and *LaNBS* were similar, mRNA accumulated in high amount in bulb and root, and low transcript levels were detected in leaf, stem, and flower (Figure 6c,d). The *NR* expression pattern was different. *LaNR* and *NpNR* mRNA specifically accumulated at high levels in bulbs, and in low amount in other tested tissues (Figure 6e,f). We further checked the expression pattern of *TR* in different tissues to compare tissue expression profiles between *NR* and *TR* and found that *NpTR* and *LaTR* genes were widely expressed in all the examined tissues (Figure S13a,b). These results suggest that the highest transcript levels of *NR* were in bulbs, which parallels with the high expression for *NBS* transcripts in the bulbs. This high expression of *NBS* and *NR* correlates with the higher AAs content in bulbs of *N.* papyraceus^15^ and *L. aestivum*^16^.

## DISCUSSION

To date, PR10/Bet v1-like proteins have been directly implicated in the biosynthesis of two alkaloid classes – the AAs and the benzylisoquinoline alkaloids (BIAs)^26,27^. Although end-product alkaloids within these two classes are structurally distinct, the pathways have similar biogenic origins. In the BIAs pathway, NCS catalyzes the condensation between dopamine and 4-HPAA to form (*S*)-norcoclaurine. The initiation of AAs’ biosynthesis is proposed to occur via condensation of tyramine and 3,4-DHBA to yield the imine norcraugsodine, then reduced to produce norbelladine (Figure 1). Our results confirm that the condensation of tyramine and 3,4-DHBA by *La* and *Np*NBS forms norbelladine at low levels, but not norcraugsodine^14,16^, despite the absence of a cofactor for the reduction in the reaction mixture. In our enzymatic assays, even the prepared standard solution of norcraugsodine was highly unstable, indicating that norcraugsodine is difficult to detect because of its instability. Still, the imine reduction of norcraugsodine intermediate into norbelladine by NBS is surprising. NBS is part of the PR10/Bet v 1-like enzymes, which are not known to use cofactors for their activity, with few reported exceptions^28^. Although unlikely, low but sufficient amount of bacterial components carrying cofactors could remain in the purified protein and help with the reduction. However, in our enzymatic assays, the addition of NADPH to the reaction did not significant increase norbelladine production by NBS. Alternatively, *Np* and *La*NBS could possess a reductase activity as found in PR-10 protein *Ca*ARP from chickpea^28^, but it is improbable as they lack the conserved motifs corresponding to catalytic signatures of short chain dehydrogenase/reductase (SDR) (Y-X_3_-K) and aldo/keto reductase (AKR) (Y-X_27–30_-K) with the common tyrosine residue^28^. On the other hand, *NpKA*NR was previously reported to also yield norbelladine following incubation with tyramine, 3,4-DHBA and NADPH, albeit at low yield^12^. We confirmed that low but detectable amounts of norbelladine are produced from 3,4-DHBA and tyramine with NR isolated for *N. papyraceus* and *L. aestivum* in enzymatic assays. In theory, other SDR members with imine reduction capability could be able to catalyze this reaction as proposed previously^20^. In our study, only NR, but not TR, the other identified SDR member, was specifically capable of catalyzing this reaction. The low yield of NBS and NR reported reactions prompted us to hypothesize that they could work together to channel the substrates rapidly and effectively for their condensation into norcraugsodine followed by an immediate reduction into norbelladine. Their interaction could help to improve their catalytic activity, avoid degradation of unstable norcraugsodine by decreasing its transit time, and prevent feedback regulation of norbelladine production.

We used homology modelling and docking study to model our hypothesis and gain insight of the ligand-enzyme interaction involved in norbelladine synthesis as a two-step reaction. Although performing docking on predicted structures has its limitation, it provides a general scheme on the possible interactions of NBS and NR with their respective ligands that is consistent with the enzymatic reactions they performed. Our proposed model of norbelladine synthesis involves NBS first where the condensation of 3,4-DHBA and tyramine subunits yields an imine/iminium intermediate (Schiff base)(*i.e.* norcraugsodine), followed by NR with a reduction to yield norbelladine (Figure 1). Similarly, to the NCS catalyzed reaction^29,30^, homology modeling and docking results suggest that NBS Tyr68, Lys83 and Glu71 surround the active site and interact with 3,4-DHBA and tyramine, probably playing key roles in norcraugsodine synthesis. In this proposed reaction, Lys83 would transfer a proton from the ammonium ion of tyramine to the carbonyl oxygen of 3,4-DHBA, stabilizing a partial positive charge on the carbon atom. Lys83-assisted nucleophilic attack of tyramine amine group to the aldehyde carbonyl would lead to the release of a water molecule from the carbinolamine moiety, generating the imine double bond, in a similar reaction to NCS’s^29^. Finally, norcraugsodine would be formed following an electrophilic attack and deprotonation assisted by the carboxyl moiety of Glu71, acting as a base. Tyr68 could contribute to the electrostatic properties of the active site and by shaping the cavity entrance. Following its synthesis, norcraugsodine would then be reduced to norbelladine inside NR active site with the assistance of NADPH. In SDR enzymes, the catalytic dyad Tyr175 and Lys179 are conserved and work together in the active site, the tyrosine serving as a general acid to protonate the substrate keto-group, while the lysine lowers the tyrosine’s pKa to promote proton donation. Noroxomaritidine reduction occurs in a similar structural and chemical arrangement: the substrate ketone is positioned in proximity to Tyr175 and NADPH^12^. Electrostatic interaction with Lys179 reduces the pKa of Tyr175 to polarize the noroxomaritidine carbonyl group for protonation and hydride transfer, to yield the ketone product with a reduced carbon-carbon double bond^12^. Similarly, to noroxomaritidine crystal structure in complex with NADPH and tyramine or piperonal, docking results showed that norcraugsodine phenol cycle binds near Phe214, its amine group is positioned close to Tyr173 and NADPH C4, and its dihydroxybenzene binds to Glu224. Lys177 in the proximity of Tyr173 could allow the tyrosine hydroxyl group to serve as a general acid with a hydride transfer from NADPH, yielding norbelladine with its reduced carbon-nitrogen double bond. Consistent with reactions catalyzed by the SDR families, docking studies suggest that NADPH, Tyr173 and Lys 177 play key roles in norcraugsodine reduction.

We tested our model pathway of norbelladine biosynthesis using several enzymatic assays. Our results confirmed that when both enzymes are present in the reaction in a single-step, or in a two-step reaction, significantly higher levels of norbelladine were produced than when using each enzyme separately. The highest yield (12-fold increase) was obtained when the assay was carried out in a two-step manner, NBS first and NR second order. These results showed that NBS and NR can cooperate to produce norbelladine *in vitro*. *In vivo*, enzymes perform their functions in specific subcellular compartments^31^. We show that NBS and NR are colocalized to the cell cytoplasm and nucleus. Their colocalization is consistent with their possible cooperation. We propose that the site of their activity is in the cytoplasm as previous studies have shown that *La* tyrosine decarboxylase 1 (TYDC1), which catalyzes the conversion of tyrosine to tyramine was localized to the cytosol^32^, as were *O*-methyltransferase 1 from *Lycoris aurea* (*O*MT) and the N4*O*MT from *Lycoris longituba,* which catalyze the *O*-methylation of norbelladine to 4’-*O*-methylnorbelladine^33^. In addition, there is some evidence of cytoplasmic localization for tyramine and 3,4-DHBA^34^. Thus, our results support the hypothesis that the early reactions of AAs are biosynthesized in the cytosol. In the Amaryllidaceae plants, we observed combined high expression of both NBS and NR in bulbs of *N. papyraceus* and *L. aestivum,* reinforcing their probable cooperation to produce norbelladine *in vivo*.

Then, we investigated the possibility that they interact together in yeast and *in planta*. First, we observed that both NBS and NR proteins physically interact as homodimers in yeast and *in planta*, suggesting that this conformation is important for their function *in vivo*. Then, our results in yeast suggest that the two enzymes directly interact with each other and that additional plant proteins are not required for this interaction. These results support the hypothesis that both NBS and NR work together in a metabolon, where NBS as a homodimer synthesizes the intermediate norcraugsodine, readily converted to norbelladine by NR, present as a homodimer in the same protein complex. The formation of metabolons allows the intermediate product to be passed directly from one enzyme into the active site of the next consecutive enzyme in the metabolic pathway^35^. It is also possible that allosteric regulation of NBS by NR (or vice-versa) could explain the enhanced production of norbelladine when both enzymes interact. NBS or NR could play a chaperone-like role in guiding the folding of the norcraugsodine intermediate or of the second enzyme, as has been previously suggested for PR10/Bet V1-like members^26,27^. Future studies will shed light on how different sequential enzymes in AAs biosynthesis forms structural-functional complex and how this metabolon is regulated.

In conclusion, our study establishes that NBS and NR cooperatively catalyze the biosynthesis of norbelladine. We show that the two enzymes localize to both the cytoplasm and nucleus, are expressed at high levels in bulbs and physically interact with each other in yeast and *in planta*. This study unravels the reactions catalyzing the first key steps involved in the biosynthesis of all AAs. Deciphering norbelladine synthesis will facilitate the development of the biosynthetic tools required to produce AAs *in vitro* and help fight human diseases, such as biosynthesize galanthamine in heterologous hosts to treat the symptoms of Alzheimer’s disease.

## EXPERIMENTAL PROCEDURES

### Plant materials and growth condition

Paperwhite narcissus (*Narcissus papyraceus*) and summer snowflake (*Leucojum aestivum*) bulbs were purchased from Vesey’s (York, PE, Canada). Bulbs were planted in plastic pots using autoclaved AGRO MIX G6 potting soil (Fafard, Saint-Bonaventure, QC, Canada). The plants were kept at room temperature with exposure to tube lighting in long day (16 h of light/8 h of dark) conditions until flowering. The plants were watered when necessary to keep the soil moist. Different tissues such as bulbs, roots, stems, leaves, and flowers were collected, flash frozen in liquid nitrogen, and stored at − 80°C until further used. *Nicotiana benthamiana*^36^ plants were grown in a growth chamber in long day (16 h of light/8 h of dark) conditions at 22°C.

### Bacterial and yeast strains and growth conditions

The bacteria used in this study were *Escherichia coli* DH5α (Invitrogen), *E. coli* Rosetta (DE3) pLysS (Novagen), and *Agrobacterium tumefaciens* GV3101^37^. The yeast strain (*Saccharomyces cerevisiae*) used is Y2HGold (Clontech Laboratories). The bacteria were grown in Luria-Bertani (LB) medium supplemented with the appropriate antibiotics at the following temperatures: *E. coli* at 37°C; and *A. tumefaciens* at 28°C. Antibiotics were used at the following concentrations (µg/mL): ampicillin, 100; chloramphenicol, 34; kanamycin, 50; rifampicin, 50; gentamicin, 30. Yeasts were grown at 30°C in selective synthetic complete medium.

### PCR amplification, cloning, and construction of vectors

RNA was extracted from *N. papyraceus* and *L. aestivum* bulbs using CTAB (cetrimonium bromide) method as described by Singh and Desgagné-Penix (2017). cDNA was synthesized from 1 µg RNA samples using SensiFAST cDNA synthesis kit (Bioline) according to manufacturer’s protocol. The open reading frame (ORF) of full length *NpNBS*, *NpNR*, *NpTR*, *LaNBS*, *LaNR*, and *LaTR* were amplified from bulbs cDNA using PrimeStar GXL premix (TaKaRa Bio) in 50 μL reaction with 0.2 μM forward and reverse primers (Supplementary Table S2). PCR program parameters: 2 min 98°C 1 cycle, 10 s 98°C, 20 s 55°C, 1 min 72°C for 35 cycles, 5 min 72°C 1 cycle, and final infinite hold at 4°C. A classical restriction digestion based cloning approach was used to clone and create all the desired vectors.

For protein expression in *E. coli*, *NpNBS*, *NpNR*, *NpTR*, *LaNBS*, *LaNR*, and *LaTR* were fused to the C-terminus of the maltose binding protein (MBP) in the pMAL-c2x vector (New England Biolabs). Precisely, the full-length ORFs were amplified from cDNA by PCR using primers reported (respective restriction enzyme sites are underlined, Supplementary Table S2). PCR products were cleaned using GenepHlow Gel/PCR kit (Geneaid). Purified PCR products of *NpNBS*, *NpNR*, *LaNBS*, and *LaNR* were digested with *Bam*HI and *Hin*dIII, while *NpTR* and *LaTR* were digested with *Bam*HI and *Sal*I and ligated into pMAL-c2x vector digested with *Bam*HI/*Hin*dIII and *Bam*HI/*Sal*I, respectively, using T_4_ DNA ligase (New England Biolabs). The recombinant plasmids were transformed into chemically competent *E. coli* DH5α cells by heat shock transformation and colonies were selected on ampicillin LB agar plates. The positive clones were identified by colony PCR in a 20 μL reaction using Taq DNA polymerase with ThermoPol buffer (New England Biolabs) with PCR parameters: 5 min 95°C 1 cycle, 30 s 95°C, 40 s 55°C, 1 min 68°C for 30 cycles, 5 min 68°C 1 cycle, and final infinite hold at 4°C. The resulting plasmids were verified by DNA sequencing to ensure the correct sequence and exclude undesired mutations.

For split luciferase complementation assays (SLCA) in *N. benthamiana* leaves, the *NpNBS*, *NpNR*, *NpTR*, *LaNBS*, *LaNR*, and *LaTR* genes were cloned into pCAMBIA1300:CLuc fused to the <u>C</u>-terminal (398-550 amino acids) of firefly <u>luc</u>iferase (CLuc), and *NpNBS*, *NpNR*, *LaNBS*, and *LaNR* into pCAMBIA1300:NLuc fused to the <u>N</u>-terminal (2-416 amino acids) of firefly <u>luc</u>iferase (NLuc) and driven by the CaMV 35S promoter^38^. The full-length ORFs were amplified by PCR using reported primers from cDNA (respective restriction enzyme sites are underlined, Supplemental Table S2). After cleaning up using GenepHlow Gel/PCR kit (Geneaid), PCR products were digested with the mentioned restriction enzymes pair and ligated into the corresponding sites of pCAMBIA1300:CLuc or pCAMBIA1300:NLuc vectors using T_4_ DNA ligase. The recombinant plasmids were transformed into chemically competent *E. coli* DH5α cells and colonies were selected on kanamycin LB agar plates. The positive clones were identified by colony PCR. The resulting binary vectors were verified by DNA sequencing.

For yeast two-hybrid assays, genes encoding full-length *NpNBS*, *NpNR*, *NpTR*, *LaNBS*, *LaNR*, and *LaTR* were amplified from *N. papyraceus* and *L. aestivum* bulbs cDNA and cloned into the pGBKT7 (bait), or pGADT7 (prey) vectors (Clontech Laboratories) in frame with the GAL4 DNA binding domain (DNA-BD) or GAL4 activation domain (AD). The full-length genes were amplified by PCR using primers reported (respective restriction enzyme sites are underlined, Supplemental Table S2). After cleaning up using GenepHlow Gel/PCR kit (Geneaid), PCR products were digested with the mentioned restriction enzymes pair and ligated into the corresponding sites of pGBKT7 (bait), or pGADT7 (prey) vectors using T_4_ DNA ligase. The recombinant plasmids were transformed into chemically competent *E. coli* DH5α cells. Colonies with bait plasmids were selected on kanamycin while colonies with prey plasmids were selected with ampicillin LB agar plates. The positive clones were identified by colony PCR. The resulting plasmids were verified by DNA sequencing.

For subcellular localization, *NpNBS*, *NpNR*, *LaNBS*, and *LaNR* coding sequences were fused upstream to the gene encoding the yellow fluorescence protein (YFP) in the pBTEX binary vector under the control of the CaMV 35S promoter^39^. The full-length ORFs were amplified (respective restriction enzyme sites are underlined, Supplemental Table S2). The PCR products were cleaned up using GenepHlow Gel/PCR kit (Geneaid), digested with *Kpn*I/*Xba*I, and ligated into pBTEX-YFP vector digested with *Kpn*I/*Xba*I using T_4_ DNA ligase. The recombinant plasmids were transformed into chemically competent *E. coli* DH5α cells and colonies were selected on kanamycin LB agar plates. The positive clones were identified by colony PCR. The resulting plasmids/binary vectors were verified by DNA sequencing.

### Expression and purification of MBP fusion proteins in *E. coli*

*NpNBS*, *NpNR*, *NpTR*, *LaNBS*, *LaNR*, and *LaTR* were cloned into the pMAL-c2x vector. The purified plasmids were transformed using heat shock transformation into chemically competent *E. coli* Rosetta (DE3) pLysS strain for protein expression. Transformed cells were selected on LB plates with ampicillin, and chloramphenicol overnight at 37°C. The positive colonies were screened by colony PCR. A PCR positive single colony was picked and grown overnight at 37°C at 220 rpm in 12.5 mL LB broth containing ampicillin, and chloramphenicol. The overnight grown pre-culture was added into fresh 250 mL LB broth containing ampicillin, and chloramphenicol and grown at 220 rpm at 37°C to an OD_600_=0.5 to 0.6. The cultures were brought to room temperature and Isopropyl-β-D-thiogalactopyranoside (IPTG) was added to a final concentration of 0.25 mM to induce protein expression. The cultures were further incubated for 20 h at 18°C at 150 rpm. Bacterial cultures were pelleted at 10,000 rpm for 15 min at 4°C, resuspended in 25 mL column binding buffer (25mM Tris-HCl [pH 7.5], 150 mM NaCl, and 1 mM EDTA) and frozen at −80°C overnight. The cultures were thawed on ice water and 1 mM phenylmethylsulfonyl fluoride [PMSF] with 0.1x protease inhibitor cocktail (Cell Signaling Technology) were added. Cultures were then lysed using a sonicator at 41% amplitude for a total of 8 min with 15 s run and 35 s cooling time in ice. The lysates were centrifuged twice at 14,000 rpm for 20 min at 4°C to pellet the cell debris and the clear supernatants were collected. Supernatants were mixed with 500 μL of amylose resin beads (New England Biolabs) (prewashed with column binding buffer and resuspended to 50% slurry) and incubated for 1 h at 4°C with constant rocking. The mixture was passed twice through the filter columns (Thermo Scientific) with gravitational flow to retain the beads with bound proteins in the column matrix. The beads were washed with gravitational flow in the columns three times with 30 mL column binding buffer. The bead slurry was transferred to microcentrifuge tube and centrifuged at 1000 rpm for 1 min and the supernatant was removed. Finally, the bound protein from the bead pellet was eluted twice (elution 1 and 2) each time in 500 μL elution buffer (15 mM maltose in column binding buffer) by centrifugation at 1000 rpm for 1 min and supernatant/elute was collected. The protein samples were flash frozen in liquid nitrogen and stored at − 80°C. Protein quantification was done using DC protein assay kit (Bio-Rad) according to the manufacturer’s instructions with bovine serum albumin (BSA) as standard, and protein samples were fractionated by 10% (v/v) SDS-PAGE and stained with Coomassie Blue.

### Protein analysis and alignment

The *in silico* protein analysis was done by DNAMAN analysis software (Lynnon BioSoft). Protein sequences of *Np*NBS, and *La*NBS were aligned with *Narcissus pseudonarcissus* ‘King Alfred’ norbelladine synthase (*NpKA*NBS; GenBank: AYV96792.1), and *Thalictrum flavum* norcoclaurine synthase (*Tf*NCS; Genbank: ACO90248.1). Similarly, *Np*NR, *La*NR, *Np*TR and *La*TR were aligned with *N. pseudonarcissus* noroxomaritidine/norcraugsodine reductase (*NpKA*NR; KU295569) using CLUSTAL W algorithm in T-Coffee software^40^ with default parameters. Sequence alignments were formatted using Boxshade program (https://embnet.vital-it.ch/software/BOX_form.html).

### Molecular homology modelling and docking

Amino acid sequences corresponding to *Np*NBS, *La*NBS, *Np*NR, *La*NR, *Np*TR, *La*TR were uploaded on Protein Homology/analogY Recognition Engine V 2.0 (Phyre2)^41^ website, I-Tasser from Zhang lab^42^ and MOE 2020.09 software (Chemical Computing Group) to model NBS and NR proteins. Following close analysis of predicted structures and comparison by superimposition with orthologs and homologs, the most consistent models were selected. *La*NR from I-Tasser, *Np*NR and *La* and *Np*TR models from Phyre2 were used, while NBS were best modeled by MOE. MOE was further used to analyze the resulting homology model conformation and prepare receptors for docking. First, modeled structures were compared to their template crystal structures in complex with their ligands downloaded from the Protein Data Bank (for NBS: norcoclaurine synthase from *Thalictrum flavum* in complex with dopamine and hydroxybenzaldehyde 2VQ5^18^, NR: noroxomaritidine/norcraugsodine reductase in complex with NADP+ and tyramine 5FF9 *NpKA*NR^12^, TR: Tropinone reductase-II complexed with NADP+ and pseudotropine 2AE2 and 5FF9^43^, aligning amino-acid sequences and then superimposing the structures.

Structure preparation consisted of correcting issues, capping, charging termini, selecting appropriate alternate, and calculate optimal hydrogen position and charges using Protonate 3D. Fixed receptor and tethered active site energy minimization was performed for each modeled protein in presence of template ligands prior to docking. Ready to dock ligands were uploaded from ZINC15^44^, when available, or manually drawn (from smiles), washed, prepared, and minimized with MOE. The MMFF94× force field was used. Receptors active site was predicted using MOE Site Finder and used as docking site to place ligand using Triangle Matcher as placement method for 200 poses, and tethered induced fit as refinement to perform flexible docking. Ten resulting poses were analyzed, and most probable poses based on literature description of templates active site, on comparison with crystalized templates interactions and on scores are presented. For NBS, 3,4-DHBA was docked first, the most consistent pose was further included in the active site used for tyramine docking. Similarly for NR, NADPH was docked first, the most consistent pose compared to template crystals was further included in the active site used to dock norcraugsodine. PLIP was used to analyze interaction between ligands and receptors^21^, further processed using PyMOL (Shrödinger).

### Substrates and standards preparation

Norbelladine and norcraugsodine were synthesized as previously described^14^. Standard solutions of 3,4-DHBA (Fisher Scientific), tyramine (Sigma-Aldrich), and papaverine (Sigma-Aldrich) were prepared at 1000 ppm in LC-MS grade methanol (Sigma-Aldrich). Standard solutions of norbelladine and norcraugsodine were prepared as previously described^14^. From these standard solutions, dilutions were performed to obtain working solutions of 100 ppm in methanol, and 1 ppm in the mobile phase (ammonium acetate 10 mM (Sigma-Aldrich) (pH 5.0), and acetonitrile (Sigma-Aldrich) [60:40]). Standard and solutions were stored in the dark at −20°C.

### Enzymatic assays

Single enzyme assays were performed at 35°C for 2 h. All the enzymatic reactions were terminated by the addition of 10 μL of 20% trichloroacetic acid (TCA). Negative controls were purified MBP-tag protein from *E. coli*, and reactions without substrate or cofactor.

The catalytic activity of the purified NBS enzymes were analyzed following the method of Singh et al., 2018^14^ with minor modifications. Reactions were conducted using 80 μg of purified protein in 100 mM HEPES buffer (pH 6.0), with 10 μM tyramine and 300 μM 3,4-DHBA, in a total volume of 100 μL. Reaction components were equilibrated at 35°C and the reaction was started by the addition of enzyme to the substrate and buffer mixture.

NR and TR single enzyme assays were performed as previously reported by Kilgore et al., 2016^12^ with minor modifications. The assay mix contained 60 μg of purified protein, 10 μM tyramine, 300 μM 3,4-DHBA, and 1 mM NADPH in 100mM sodium citrate buffer (pH 6.0), in a total volume of 100 μL.

The assays with NBS and NR or TR together in a single-step reaction contained 80 μg of purified NBS enzyme, 60 μg of purified NR/TR enzyme, 10 μM tyramine, 300 μM 3,4-DHBA, and 1 mM NADPH in 100mM HEPES buffer (pH 6.0), in a total volume of 100 μL.

The assays with NBS and NR/TR in two-step reactions were conducted as follows: 80 μg of purified NBS enzyme in 100 mM HEPES buffer (pH 6.0), with 10 μM tyramine and 300 μM 3,4-DHBA, in a total volume of 100 μL was incubated at 35°C for 2 h. The NBS enzyme was inactivated by boiling at 95°C for 10 min, sample was centrifuged, and the supernatant (100 μL) was used as norcraugsodine solution, and 60 μg of purified NR/TR enzyme and 1 mM NADPH was added and incubated for additional 2 h.

Following the reaction termination, papaverine (1000 PPM) was added to all the reaction serving as an internal standard for the relative quantification of the concentration of detected compounds. All reactions were performed in triplicates. The reaction samples were diluted 100x with mobile phase (10 mM ammonium acetate [pH 5.0], and acetonitrile [60:40]) and analysis of the enzymatic product (norbelladine) using a high-performance liquid chromatography (HPLC) system coupled with a tandem mass spectrometer (MS/MS) was carried out as described by (Tousignant *et al.*, 2022)^16^.

### *Agrobacterium*-mediated transient expression

The YFP- and LUC-fusion binary vectors were transformed into *A. tumefaciens* strain GV3101 by electroporation and colonies were selected on LB agar plates with rifampicin, kanamycin, and gentamicin at 28°C. The positive colonies were confirmed by colony PCR using PCR parameters: 10 min 95°C 1 cycle, 30 s 95°C, 40 s 55°C, 1 min 68°C for 30 cycles, 5 min 68°C 1 cycle, and a final infinite hold at 4°C. For transient expression, cultures of *A. tumefaciens* were grown overnight in 5 mL LB broth with rifampicin, kanamycin, and gentamicin at 28°C. The cultures were pelleted at 8000 rpm for 5 minutes at room temperature, washed three times with 10 mM MgCl_2_, resuspended in 5 mL of induction medium (10 mM MgCl_2_, 10 mM MES [pH 5.6], and 200 µM acetosyringone), and incubated at 28°C with shaking at 200 rpm for 3-4 h. *A. tumefaciens* cultures were diluted in the induction media to OD_600_=0.25 and infiltrated into young but fully expanded leaves of five-week-*old N. benthamiana* plants using a 1 mL needleless syringe. After agroinfiltration the plants were incubated in a growth chamber in long day (16 h of light/8 h of dark) conditions at 22°C for 48 h until leaf discs were harvested.

### Split luciferase complementation assay

*NpNBS*, *NpNR*, *NpTR*, *LaNBS*, *LaNR*, and *LaTR* genes were cloned in frame to firefly luciferase fragments in the binary vector pCAMBIA1300:NLuc or pCAMBIA1300:CLuc. The obtained binary vectors were transformed into *A. tumefaciens*. The desired NLuc- and CLuc-fusion *A. tumefaciens* combinations were mixed at 1:1 ratio (OD_600_=0.25) and coexpressed in *N. benthamiana* leaves. Split luciferase complementation assays were performed as described by (Chen *et al.*, 2008)^38^ with minor modifications. Three millimeter-diameter leaf discs were harvested at 48 h after agroinfiltration and floated abaxial side up in 100 µL of degassed water on a white 96-well plate. Samples were supplemented with 1 mM D-luciferin (Sigma-Aldrich) and incubated in the dark for 2 min with gentle shaking and additional 8 min at rest to quench fluorescence. Luminescence was measured using a Synergy H1 Microplate reader (BioTek) with integration time set to 2 s and imaged using a Gel Doc XR system (Bio-Rad).

### Yeast two-hybrid analysis

*NpNBS*, *NpNR*, *NpTR*, *LaNBS*, *LaNR*, and *LaTR* genes were either fused to the GAL4 DNA binding domain (DNA-BD) in the bait vector pGBKT7 or were fused to the GAL4 activation domain (AD) in the prey vector pGADT7 (Clontech Laboratories). The yeast strain Y2H Gold (Clontech Laboratories) was first transformed with the bait vectors (*i.e., NBS*, *NR*, and *TR* in the pGBKT7 vector) and subsequently with prey vectors (*i.e., NBS*, *NR*, and *TR* in the pGADT7 vector). The transformants were selected on synthetically defined (SD) medium lacking leucine and tryptophan (SD-LW). The interactions were verified by testing the activation of the *HIS3*, *ADE2* and *AUR1-C* reporter genes on selective media plates lacking histidine and adenine (SD-LWHA) or containing the antibiotic Aureobasidin A (AbA), respectively.

### Protein extraction

For protein extraction from *N. benthamiana* leaves, five leaf discs (1 cm diameter) were frozen in liquid nitrogen, homogenized in 300 µL extraction buffer (100 mM Tris [pH 7.5], 1% [v/v] Triton X-100, 1 mM PMSF, and 0.1x protease inhibitor cocktail), and centrifuged at 17,000g for 30 min at 4°C. The clear supernatant was collected, and protein concentration was determined using DC protein assay kit (Bio-Rad) according to the manufacturer’s instructions.

For protein extraction from yeast, 5 mL overnight-grown cultures were pelleted at 12,000g for 5 min at 4°C, resuspended in 250 µL ice-cold lysis buffer (4% [v/v] 5 N NaOH and 0.5% [v/v] β-mercaptoethanol), and incubated with 1x SDS sample buffer (30% [v/v] glycerol, 15% [v/v] β-mercaptoethanol, 37.5% [v/v] 500 mM Tris-HCl [pH 6.8], 0.15% [w/v] SDS, and a few grains of Bromophenol Blue) for 10 min at 95°C.

### Western blotting

Equal amounts of protein (100 µg) were fractionated by 10% (v/v) SDS-PAGE. Proteins from gels were transferred onto Polyvinylidene difluoride (PVDF) membrane using Trans-Blot Turbo transfer system (Bio-Rad). The membrane was equilibrated with Tris-buffered saline (TBS) buffer (20 mM Tris, 150 mM NaCl pH 7.6) for 15 min, followed by blocking of membrane for 2 h with TBS buffer containing 0.1% tween 20 (TBST), and 5% skim milk. The PVDF membrane was incubated overnight at 4°C in TBST with 5% milk containing 1:1000 dilution of specific primary antibodies. The primary antibodies used in this study are rabbit anti full-length firefly luciferase antibodies (Sigma-Aldrich), which react with both the N-terminal and C-terminal firefly LUC fragments, mouse anti-GFP/CFP/YFP monoclonal antibody (Cedarlane labs), mouse anti-Myc/c-Myc monoclonal antibody (Santa Cruz Biotechnology), and mouse anti-HA-tag monoclonal antibody (GenScript). After primary antibody incubation, the membrane was washed three times each for 5 min in TBST buffer and incubated for 30 min in TBST containing 5% skim milk and goat anti-rabbit horse radish peroxidase (GAR)-HRP or goat anti-mouse horse radish peroxidase (GAM)-HRP conjugate in 1:10,000 dilutions. The immunoblot was washed three times for 5 min each in TBST buffer and developed using clarity Western ECL substrate (Bio-Rad). Finally, the membrane was washed twice with TBST and stained with Ponceau S stain [0.5% (w/v) Ponceau S (Sigma-Aldrich) in 1% (v/v) acetic acid] for 1 min and photographed using Gel Doc XR system (Bio-Rad).

### Subcellular localization

To visualize *Np*NBS, *Np*NR, *La*NBS, and *La*NR subcellular localization, the YFP fusion proteins were expressed via *A. tumefaciens* in leaves of 5-week-old *N. benthamiana* plants. Forty-eight hours post infiltration, the abaxial epidermis of leaf discs were placed on a microscopic slide in a water drop, covered by a cover slip, and imaged immediately. Protein localization was visualized by a confocal laser scanning microscope (Leica TCS SP8; Leica Microsystems) with a 40X/1.30 oil immersion objective. Images were first processed with Las AF Lite software (Leica Microsystems). CFP was used as a control for colocalization (Kruse et al., 2010). YFP was excited with an argon laser at 488 nm, while CFP was excited with a diode laser at 405 nm. Emission was detected with a spectral detector set between 500 and 525 nm for YFP and between 420 and 490 nm for CFP. Chlorophyll autofluorescence was observed with an excitation wavelength of 552 nm and the emission of fluorescence signals were detected from 630 to 670 nm. The combined images were generated using the Las X software (Leica Microsystems).

### RNA extraction and Real-time quantitative PCR

Total RNA was isolated from bulbs, roots, stems, leaves, and flowers using the TRIzol reagent (Invitrogen). Briefly, 100 mg of tissues were frozen in liquid nitrogen, fully ground, and homogenized in 1 mL of TRIzol using a mortar and pestle. The liquid was transferred to a microcentrifuge tube, incubated 5 min at room temperature and extracted with 200 µL chloroform. Following centrifugation at 12,000g for 15 min at 4°C, the upper phase containing RNA was transferred to a fresh tube. The RNA was precipitated with 500 µL of isopropanol for 10 min at room temperature and centrifuged at 12,000g for 10 min at 4°C. The RNA pellet was washed twice with 1 mL of 75% ethanol (with DEPC water) and centrifuged at 7500g for 5 min at 4°C. Finally, the RNA pellet was air dried and suspended in 40 µL of DEPC-treated water. The quality and quantity of RNA extracted from different tissues were verified on NanoPhotometer (Implen) and 1.5% (w/v) agarose gel electrophoresis. RNA samples (1 µg) were reverse transcribed using SensiFAST cDNA synthesis kit (Bioline) according to manufacturer’s protocol and subjected to Real-time quantitative PCR (RT-qPCR) using gene-specific primers (Supplementary Table S2). The experiments were performed in triple technical replicates of each plant sample. A total reaction volume of 20 μL containing 1x SensiFAST SYBR Lo-ROX mix (Bioline), 200 μM of each forward and reverse primer, and 2 µL of template cDNA (50 ng/µL) was used for RT-qPCR analysis. RT-qPCR was performed on CFX Connect Real-Time PCR System (Bio-Rad). Amplification conditions were 95°C for 3 min 1 cycle, 95°C for 10 s, and 60°C for 30 s for 40 cycles followed by dissociation step 95°C for 10 s, 65°C for 5 s and 95°C for 5 s. *LaGAPDH* and *NpHISTONE* were used as internal reference genes for *La* and *Np*, respectively. To verify the specificity of the primers, a melting-curve analysis was also performed. The threshold cycle (C_T_) value of each gene was normalized against the C_T_ value of the reference genes. Mean C_T_ values calculated from the technical triplicates were used for quantification of relative gene expression involving the comparative C_T_ method^45^. The results were analyzed, and the statistical error was calculated using CFX Maestro software (Bio-Rad).

### Accession numbers

Sequence data from this article can be found in GenBank under the following accession numbers: *N. papyraceus* norbelladine synthase (*NpNBS*; MZ054104), *N. papyraceus* noroxomaritidine/norcraugsodine reductase (*NpNR*; MF979872), *N. papyraceus* histone (*NpHistone*; MF979875), *L. aestivum* norbelladine synthase (*LaNBS*; MW971977), *L. aestivum* noroxomaritidine/norcraugsodine reductase (*LaNR*; MW971981), *L. aestivum* glyceraldehyde-3-phosphate dehydrogenase (*LaGAPDH*; MW971984), *Narcissus pseudonarcissus* ‘King Alfred’ norbelladine synthase (*NpKANBS*; AYV96792), *N. pseudonarcissus* noroxomaritidine/norcraugsodine reductase (*NpKANR*; KU295569).

## ACKNOWLEDGEMENTS

We gratefully thank Professor Hugo Germain for helpful discussion, and for providing the *N. benthamiana* seeds and other lab materials used in this study. We gratefully thank Professor Guido Sessa for providing the pBTEX, pCAMBIA1300:CLuc, and NLuc vectors and the *Agrobacterium* GV3101 strain used in this study. We thank Fatma Meddeb for timely help in obtaining the lab materials and helpful discussions. We also thank Melodie B. Plourde for helping on confocal imaging.

This work was funded by the Natural Sciences and Engineering Research Council of Canada (NSERC) award number RGPIN-2021-03218 (Discovery) to I.D-P. This work was also supported by the Canada Research Chair on plant specialized metabolism Award No 950-232164 to I.D-P. Thanks are extended to the Canadian taxpayers and to the Canadian government for supporting the Canada Research Chairs Program.

