## Supplementary Figures and Tables for "Characterization of norbelladine synthase and noroxomaritidine/norcraugsodine reductase reveals a novel catalytic route for the biosynthesis of Amaryllidaceae alkaloids including the Alzheimer’s drug galanthamine"

Figure S1

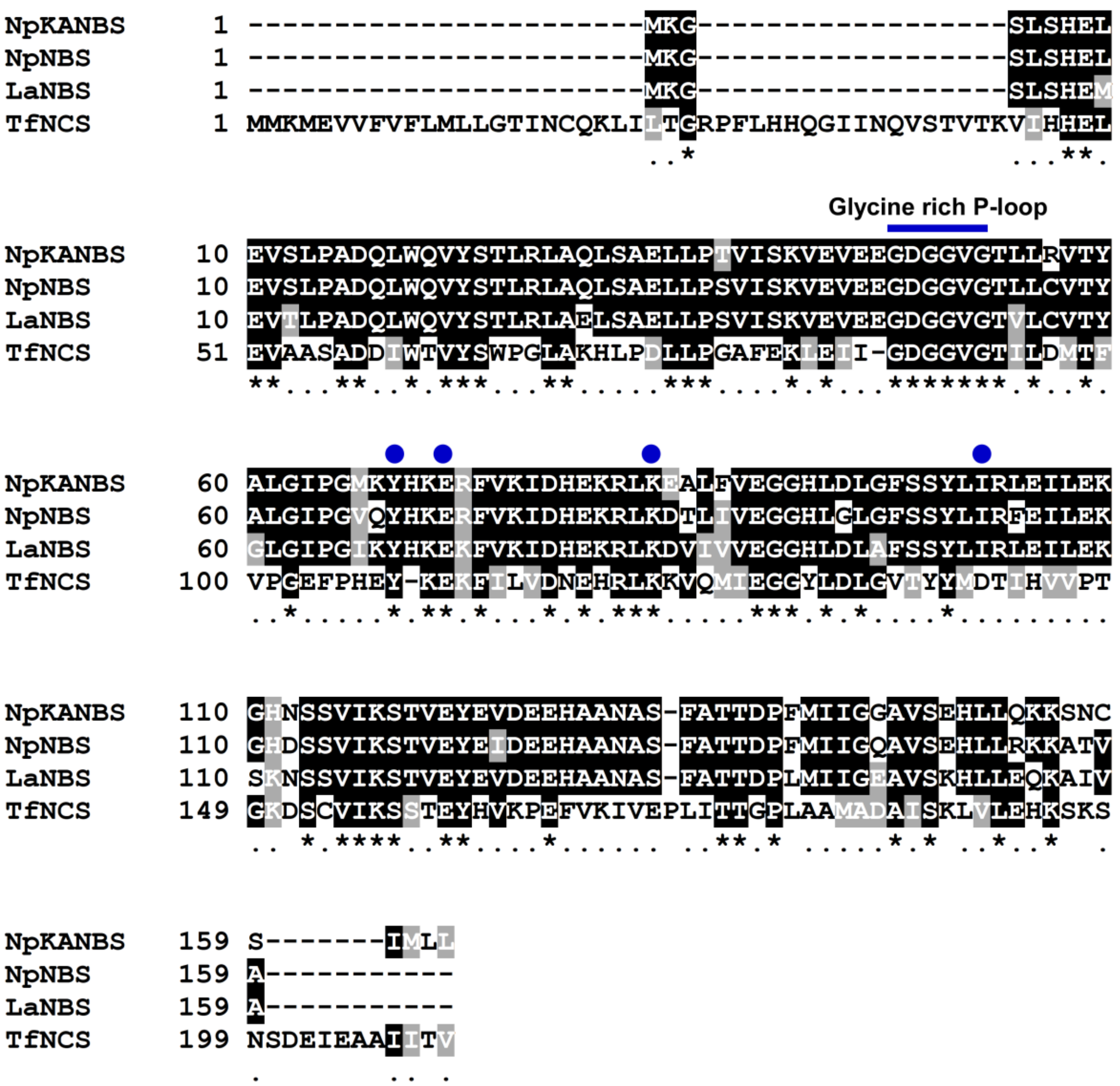


**Figure S1.** Multiple sequence alignment from the amino acid sequences of *NpKA*NBS, *Np*NBS, *La*NBS and *Tf*NCS using the CLUSTAL W algorithm. The blue bar depicts a glycine-rich P-loop domain conserved in NCS and NBS protein. Blue circles represent the Tyr, Glu, Lys and Ile residues that form the catalytic site of NBS from different species. Identical residues are marked with an asterisk (*). The positions with conservation between amino acid residues of similar properties are marked with a period (.) and numbers on the left indicate the residues position in the sequence.

Figure S2

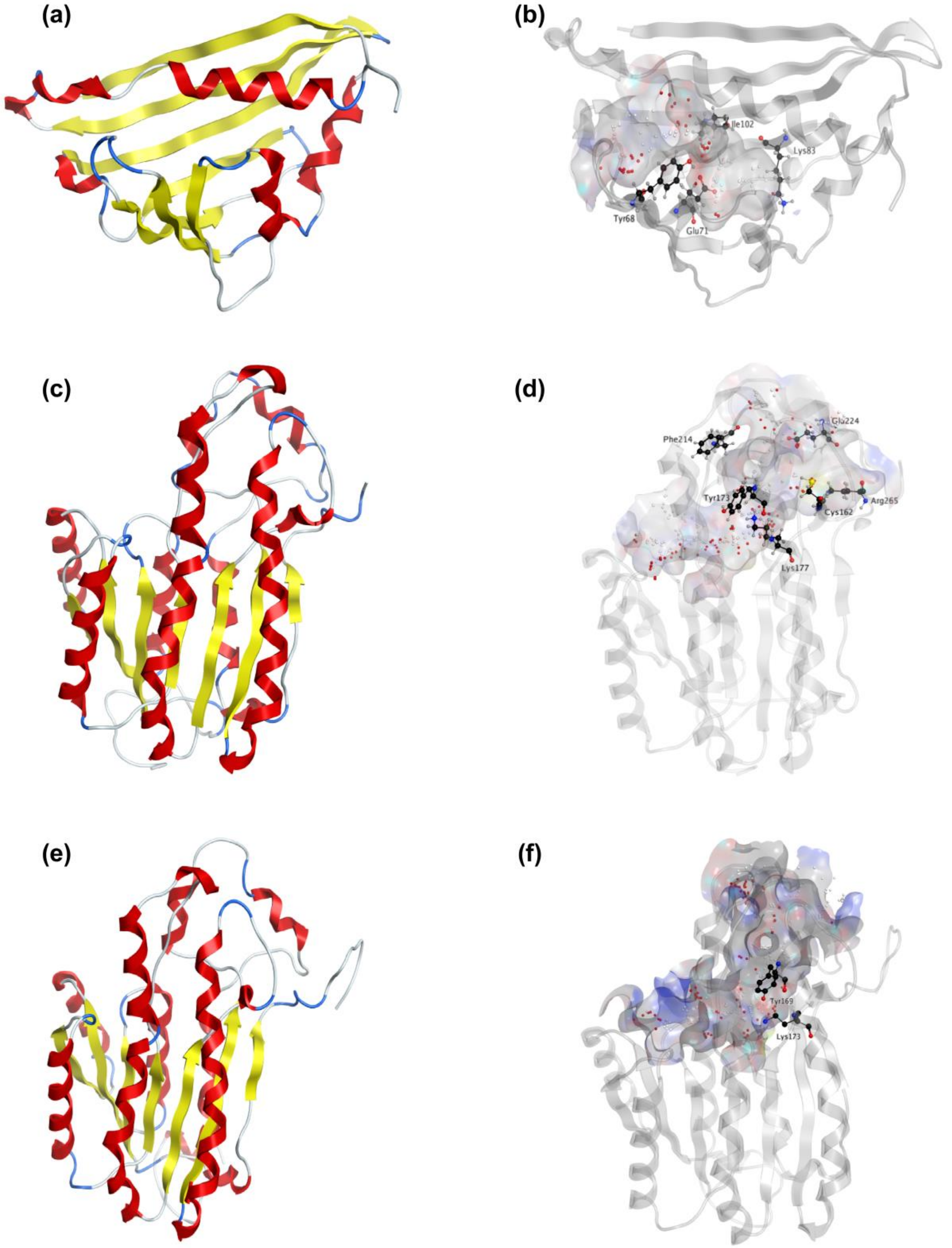


**Figure S2.** Homology modeling of *La*NBS, *La*NR and *La*TR. (a) Ribbon representation of *La*NBS with colored secondary structures. β-sheets are displayed yellow, α-helices red, loops blue. (b) Ribbon representation of *La*NBS protein with predicted active site pocket in transparent surface. The predicted ligands site computed by the Site Finder tool of MOE software is displayed as white and red alpha sphere centers inside the cavity. Conserved active site residues Tyr68 (Tyr108 from *Tf*NCS), Glu71 (Glu111), Lys83 (Lys123), and Ile102 (replacing Asp142 from *Tf*NCS) are shown as black sticks. (c) Ribbon representation of *La*NR with highlighted secondary structures. β-sheets are displayed yellow, α-helices red and loops in blue. (d) Ribbon representation with transparent surface view of predicted active site forming a catalytic tunnel that crosses the enzyme. The predicted ligands site is displayed as white and red alpha sphere centers inside the pocket. Conserved active site residues Cys162, Tyr173, Lys177, Phe214, Glu224, Arg265 surrounding the tunnel are shown as black sticks. (e) Ribbon representation of *La*TR with colored secondary structures. β-strands are displayed yellow, α-helices red, and loops blue. (f) Ribbon representation with transparent surface view of predicted active site tunnel crossing the enzyme. Predicted ligands site is displayed as white and red alpha sphere centers. Conserved catalytic Tyr169 and Lys173 are shown as black sticks.

**Figure S3**

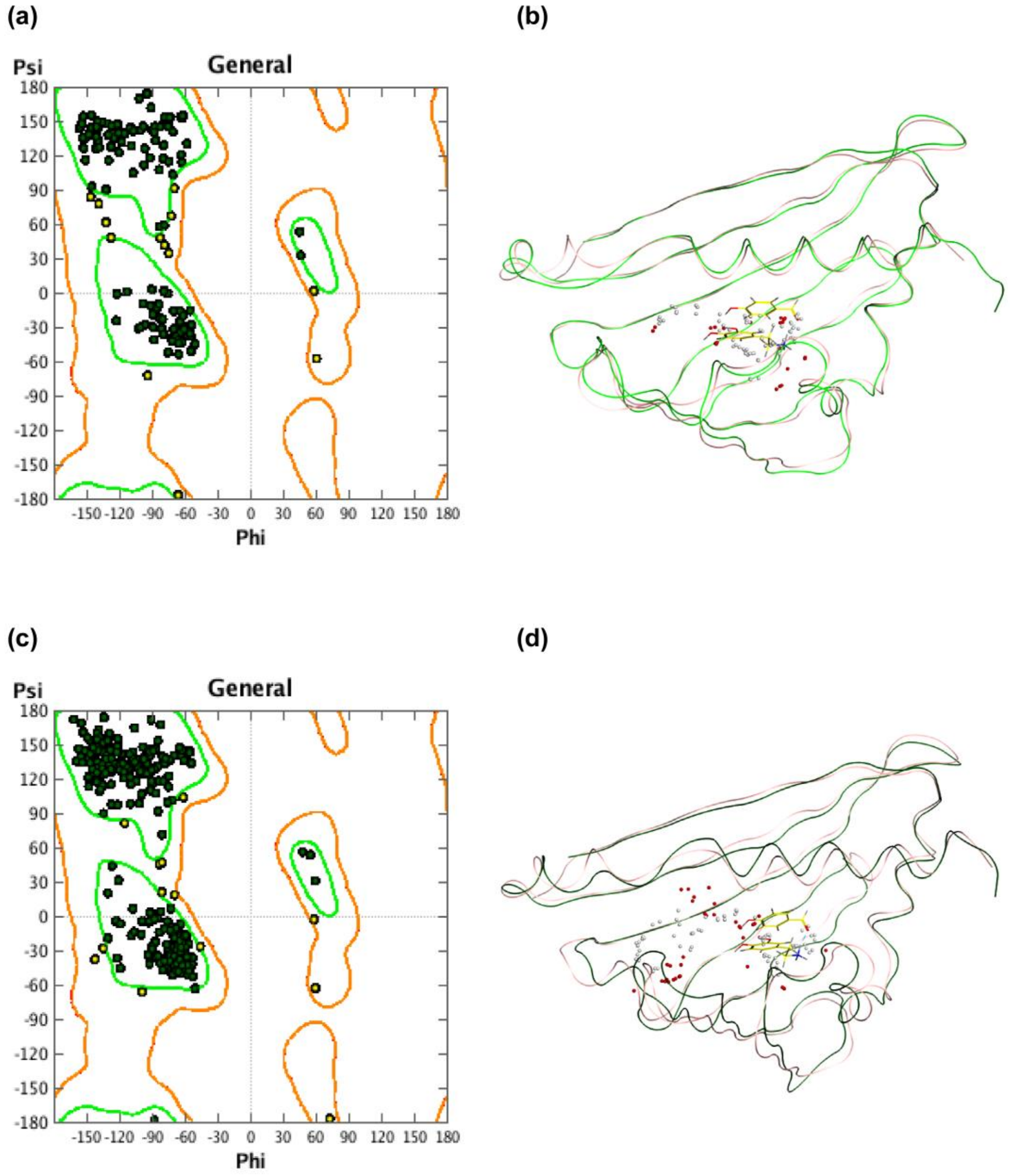


**Figure S3.** Validation of *Np* and *La*NBS modeled 3D structures. (a) Ramachandran plot of residues Psi and Phi dihedral angles of *Np*NBS homology model. Thirteen of 159 residues (8.18%, in yellow) presented allowed angles configuration while the rest had a favored geometry. *Tf*NCS (2VQ5) was used as template for homology modeling. (b) *Np*NBS thin ribbon structure (green) was superposed to *Tf*NCS 2VQ5 chain B (pink) with 39% of identity. Predicted *Np*NBS ligands site are shown as red and white alpha spheres while *Tf*NCS dopamine and hydroxybenzaldehyde ligands are displayed as thin yellow sticks. (c) Ramachandran plot of *La*NBS homology model residues Psi and Phi dihedral angles. Thirteen of 155 residues (8.39%, in yellow) presented allowed angles configuration while the rest had a favored geometry. *Tf*NCS (2VQ5) was used as template for homology modeling. (d) *La*NBS thin ribbon structure (dark green) was superposed to *Tf*NCS 2VQ5 chain B (light pink) with 41% of identity. Predicted *La*NBS ligands site are shown as red and white alpha spheres while *Tf*NCS dopamine and hydroxybenzaldehyde ligands are displayed as thin yellow sticks.

**Figure S4**

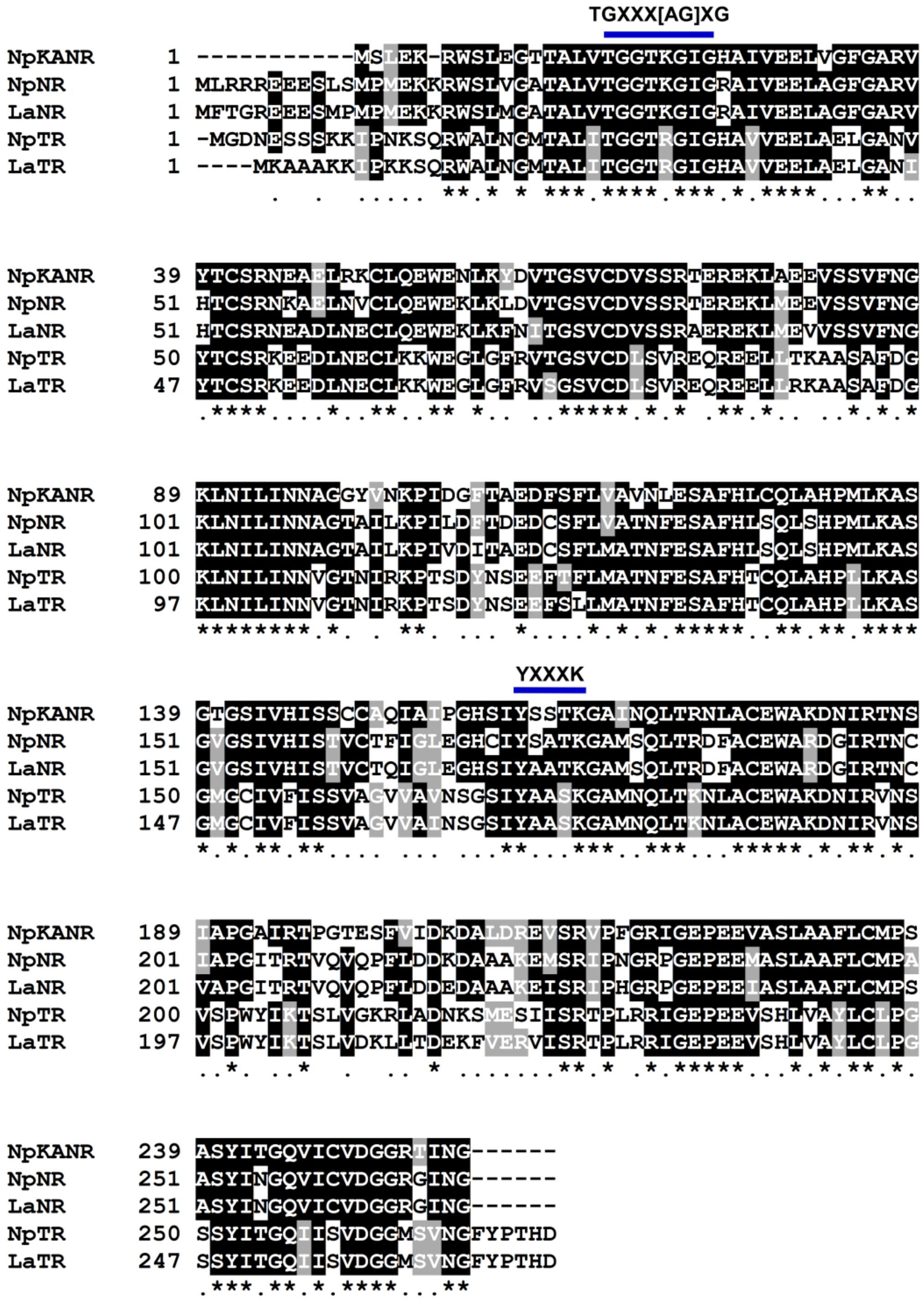


**Figure S4** Multiple sequence alignment from the amino acid sequences of *NpKA*NR, *Np*NR, *La*NR, *Np*TR and *La*TR using CLUSTAL W algorithm. The blue bars depict the TGXXX[AG]XG cofactor binding motif and a YXXXK active site motif conserved in all SDR family protein. Identical residues are marked with an asterisk (*). The positions with conservation between amino acid residues of similar properties are marked with a period (.) and numbers on the left indicate the residues position in the sequence.

**Figure S5**

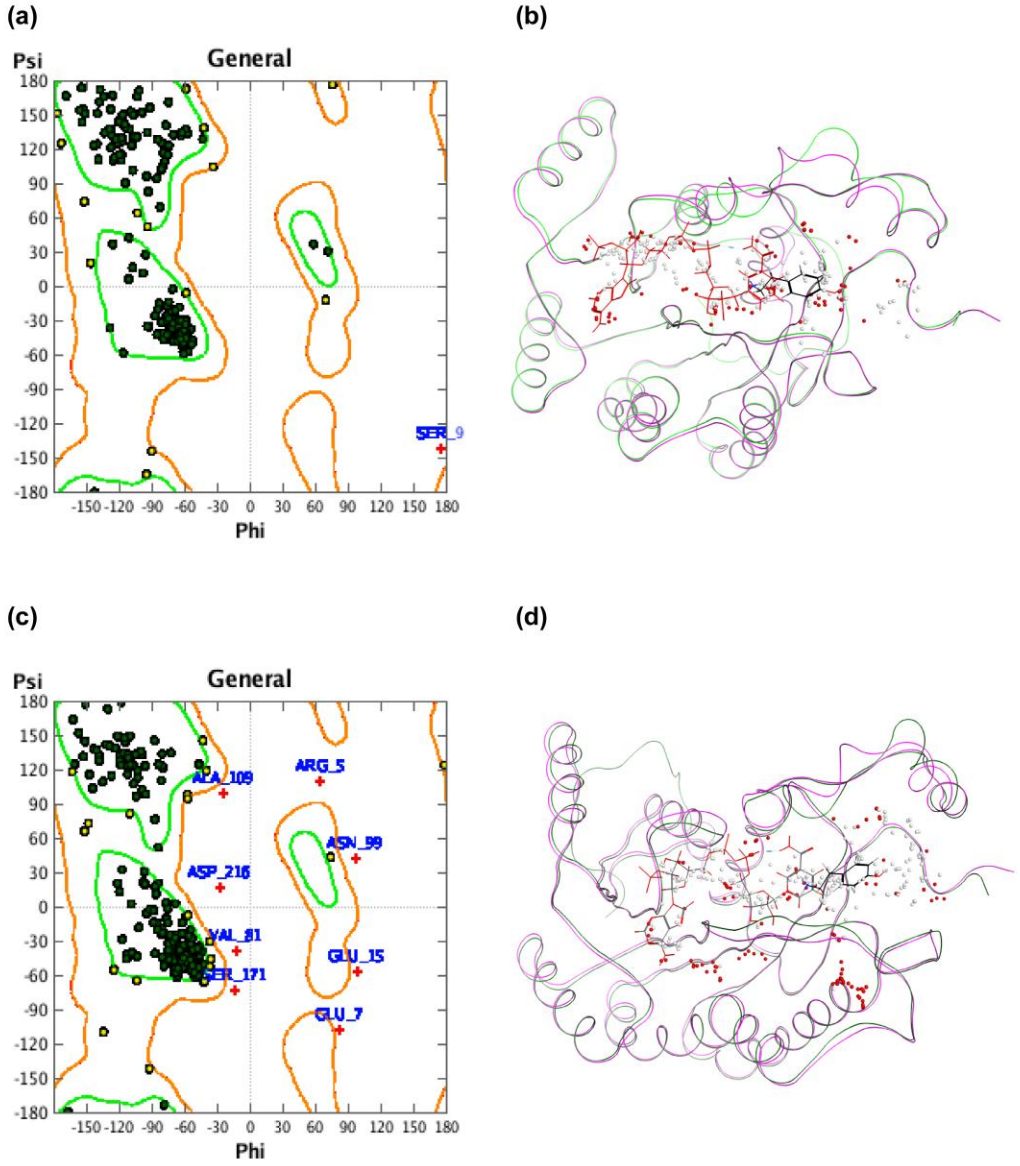


**Figure S5.** Validation of *Np* and *La*NR modeled 3D structure. 5FF9 noroxomaritidine reductase in complex with NADP+ and tyramine was used as template to model the structure of the enzyme. (a) Ramachandran plot of residues Psi and Phi dihedral angles. The graph displays 1 outlier of 269 (Ser9 in red), 12 of 269 residues (4.5%, in yellow) with allowed angles configuration while the rest had favored geometry. 252 residues (94%) modelled at >90% accuracy. (b) Active site facing view of *Np*NR ribbon structure (green) superposed to noroxomaritidine 5FF9 chain B (pink) with 75% of identity. Predicted *Np*NR ligands site are shown as white and red alpha center spheres while NADP+ and tyramine 5FF9 ligands are displayed in thin red and black sticks, respectively. (c) Ramachandran plot of residues Psi and Phi dihedral angles. The graph displays 8 outliers of 269 (3% in red), 19 of 269 residues (7.1%, in yellow) with allowed angles configuration while the rest had favored geometry. (d) Active site facing view of *La*NR ribbon structure (green) was superposed to noroxomaritidine 5FF9 chain B (pink) with 73% of identity. Predicted *La*NR ligands site are shown as white and red alpha center spheres while NADP+ and tyramine 5FF9 ligands are displayed in thin red and black sticks, respectively.

**Figure S6**

(a)

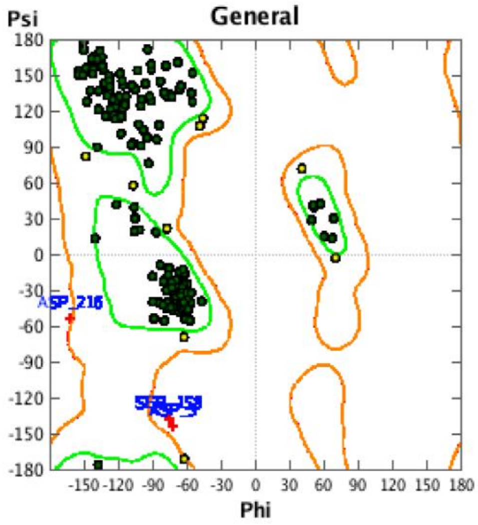


(b)

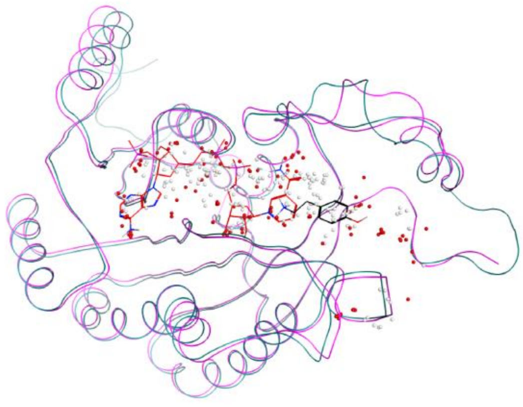

(c)

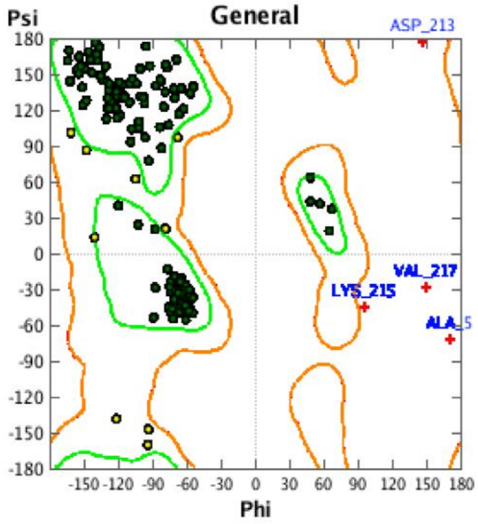


(d)

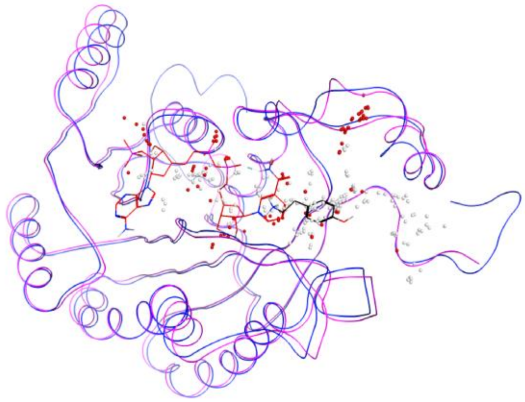

**Figure S6.** Validation of *Np* and *La*TR modeled 3D structure. (a) Ramachandran plot of *Np*TR residues Psi and Phi dihedral angles. 3 outliers of 274 (1.1%, in red), 9 of 274 residues (3.3%, in yellow) presented allowed angles configuration while the rest had favored geometry. Three templates were selected during Phyre2 modeling of *Np*TR based on heuristics to maximize confidence, percentage identity and alignment coverage. Templates are 1AE1 (56% of identity), 5FF9 (60%) and 2AE2 (56%). 18 residues at the N-terminal were modelled by *ab initio*. (b) Superposition of *Np*TR (blue) and 5FF9 (pink) predicted secondary structures. (c) Ramachandran plot of residues Psi and Phi dihedral angles. 4 outliers of 271 (1.5%, in red), 9 of 271 residues (3.3%, in yellow) presented allowed angles configuration while the rest had favored geometry. Three templates were selected during Phyre2 modeling of *La*TR based on heuristics to maximize confidence, percentage identity and alignment coverage. Templates are 1AE1 (56% of identity), 5FF9 (60%) and 2AE2 (56%). 18 residues at the N-terminal were modelled by *ab initio*. (d) **S**uperposition of *Np*TR (blue) and 5FF9 (pink) predicted secondary structures.

**Figure S7**

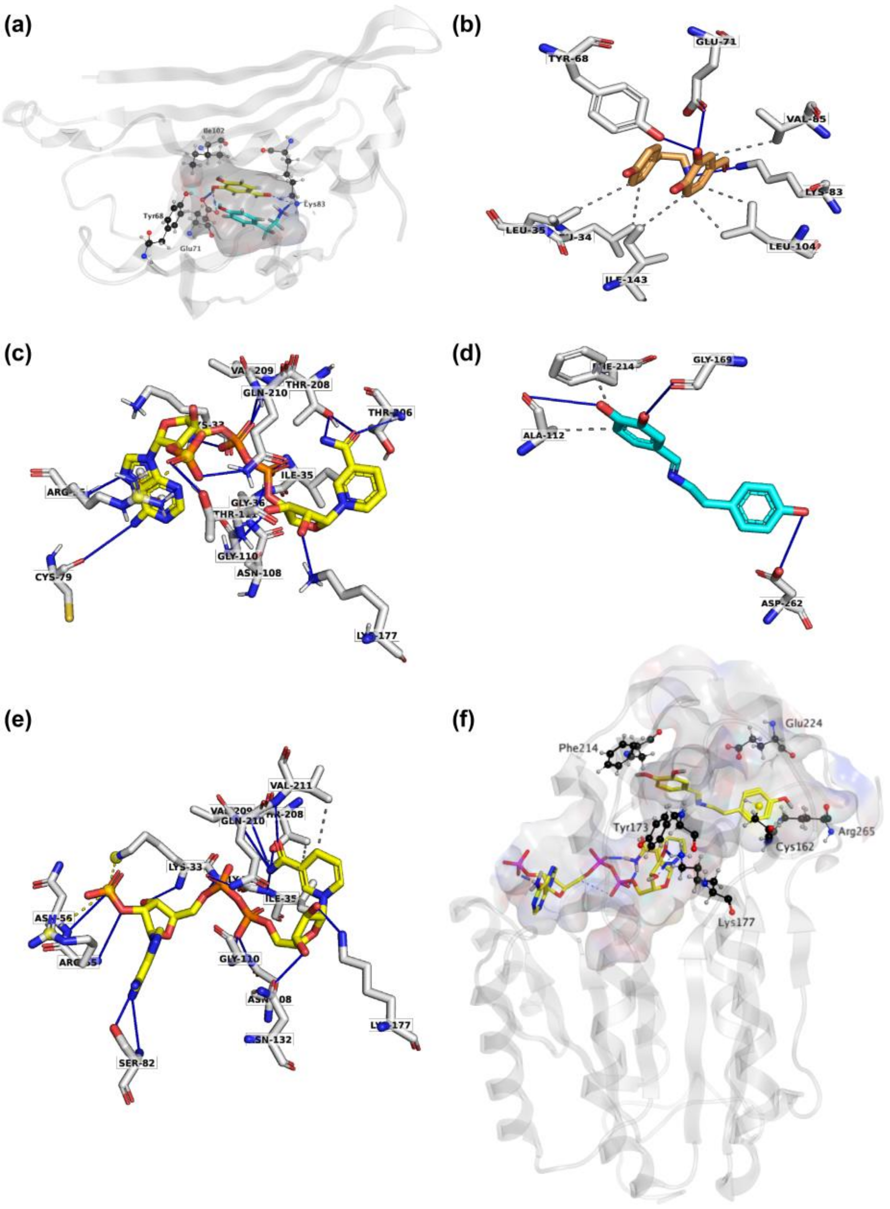


**Figure S7.** *La*NBS modeling and predicted interaction with docked 3,4-DHBAOP and tyramine. (a) Ribbon representation of *La*NBS with transparent surface-active site and black sticks catalytic residues Lys83, Tyr68, Glu71 and Ile102. Docked tyramine (turquoise) and 3,4-DHBAOP (yellow) are shown as sticks with hydrogen bonds in blue. (b) Stick representation of PLIP predicted interactions of *La*NBS (grey) with 3,4-DHBAOP (right) and tyramine (left). (c) Stick representation of PLIP predicted interactions of *Np*NR (grey) with NADPH (yellow). (d) Stick representation of PLIP predicted interactions of *La*NR (grey) with norcraugsodine (turquoise). (e) Stick representation of PLIP predicted interactions of *La*NR (grey) with NADPH (yellow). (f) *La*NR docked with NADPH and norcraugsodine. Ligands NADPH (left) and norcraugsodine (right) are represented as thick yellow sticks. Conserved active site residues Cys162, Tyr173, Lys177, Phe214, Glu224 and Arg265 shaping the binding site are shown as thin black sticks.

**Figure S8**

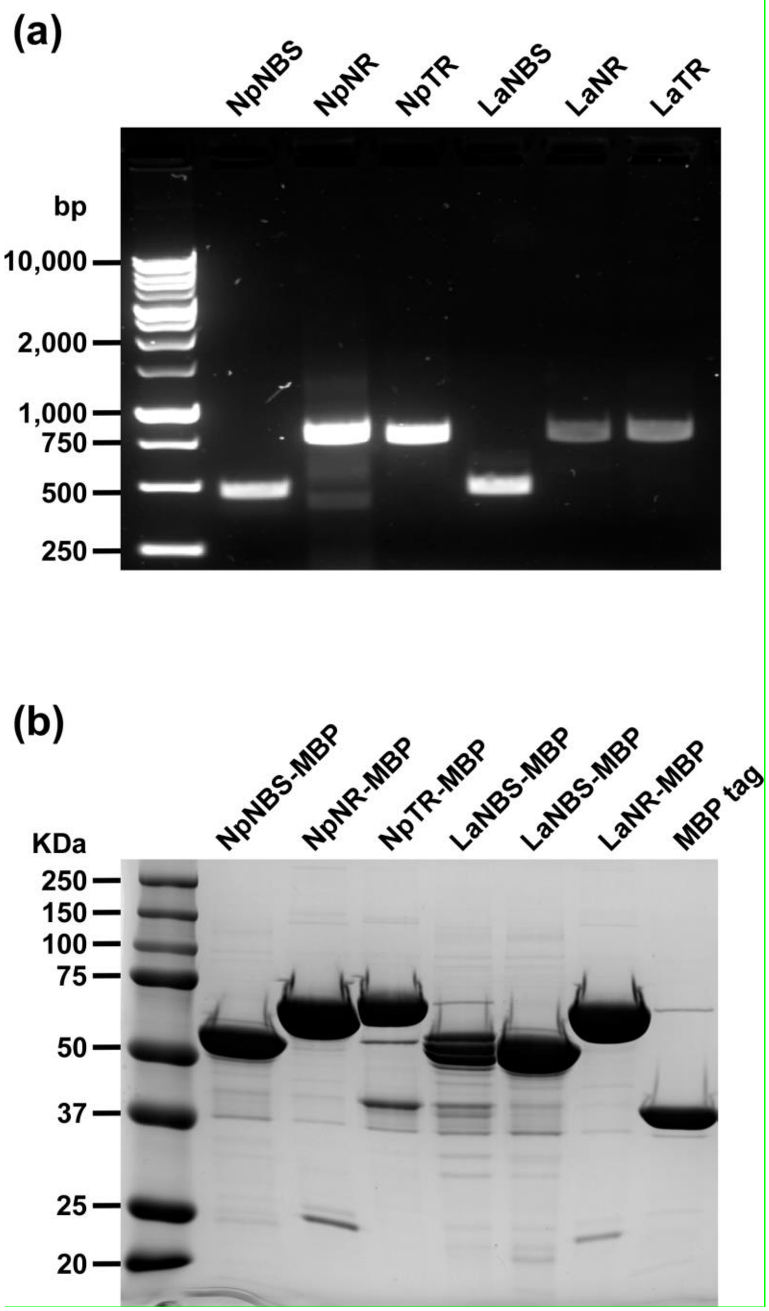


**Figure S8.** Cloning and heterologous expression of *NBS*, *NR*, and *TR* from *N. papyraceus* and *L. aestivum* in *E. coli*. (a) Agarose gel electrophoresis of PCR amplification of *NpNBS*, *NpNR*, *NpTR*, *LaNBS*, *LaNR*, and *LaTR* ORF from *N. papyraceus* and *L. aestivum* bulb cDNA library. The amplicon size is *NpNBS*, 480 bp; *NpNR*, 810 bp; *NpTR*, 825 bp; *LaNBS*, 480 bp; *LaNR*, 810 bp; *LaTR*, 816 bp. The sizes of the 1 kb ladder are positioned on the left. (b) SDS-PAGE analysis of *Np*NBS-MBP, *Np*NR-MBP, *Np*TR-MBP, *La*NBS-MBP (in duplicate), *La*NR-MBP, and MBP-tag alone purified fusion proteins. Fusion proteins were extracted from 0.25 mM IPTG induced *E. coli* Rosetta (DE3) pLysS host strain at 18°C for 20 h and purified proteins were eluted with 15 mM maltose. Numbers on the left refer to the location of standard protein molecular weight markers in kDa.

**Figure S9**

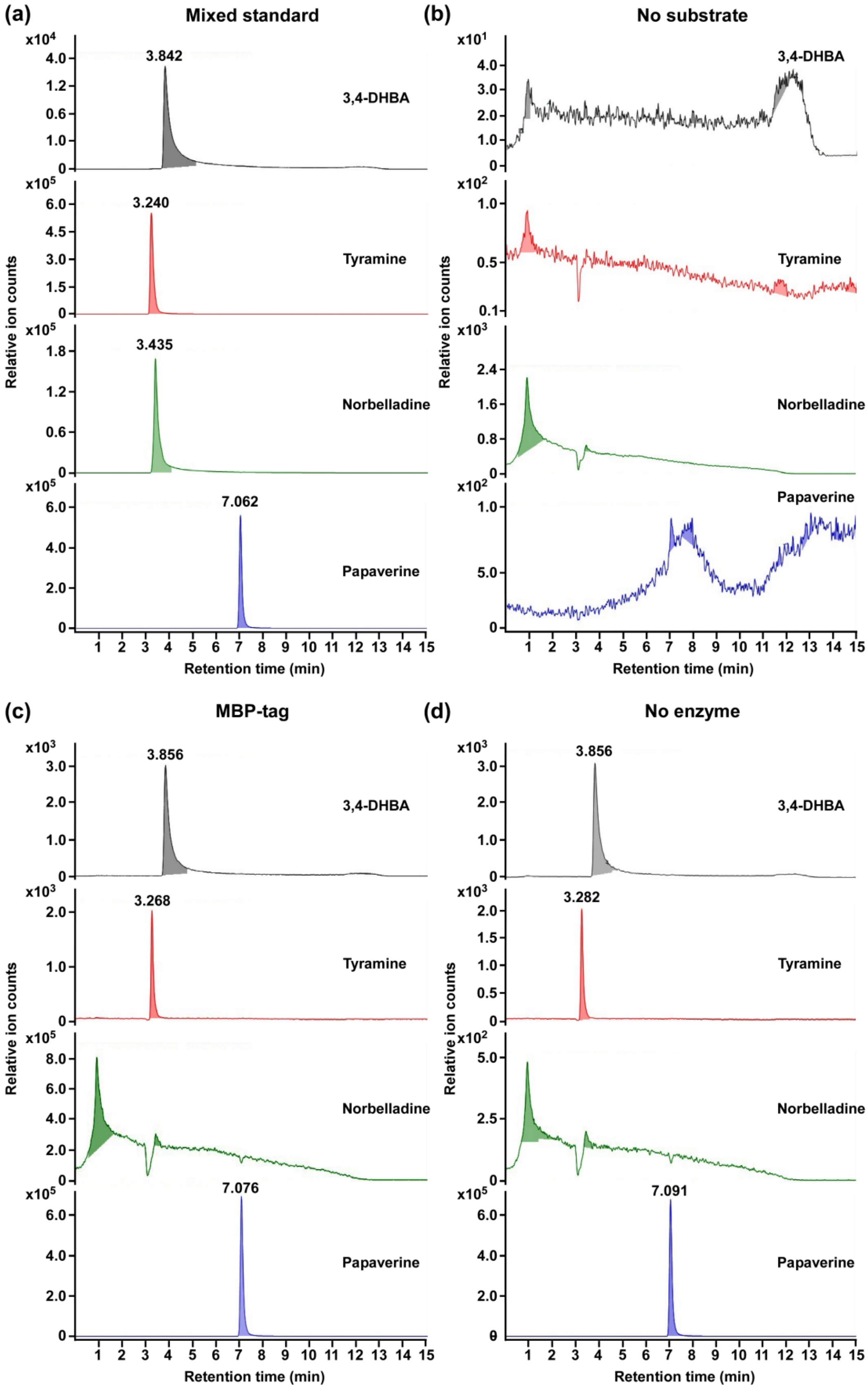
(a)
Mixed standard
(b)
No substrate
(c)
MBP-tag
(d)
No enzyme
3,4-DHBA
Tyramine
Norbelladine
Papaverine
3.842
3.240
3.435
7.062
3.856
3.268
7.076
3.856
3.282
7.091
Relative ion counts
Retention time (min)

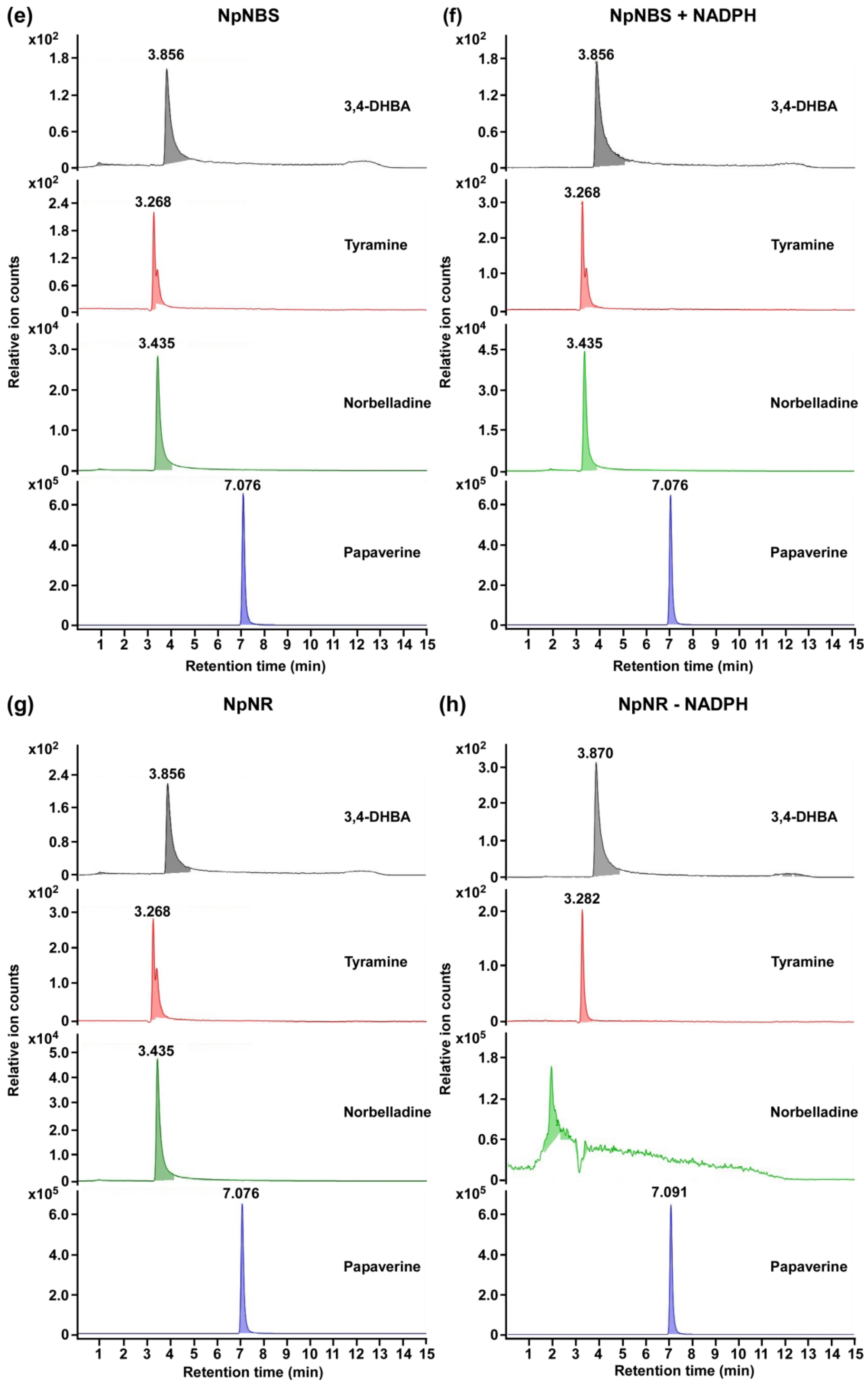
(e)
NpNBS
(f)
NpNBS + NADPH
(g)
NpNR
(h)
NpNR - NADPH
3,4-DHBA
Tyramine
Norbelladine
Papaverine
3.856
3.268
3.435
7.076
3.870
3.282
7.091
Relative ion counts
Retention time (min)
x10^2
x10^4
x10^5

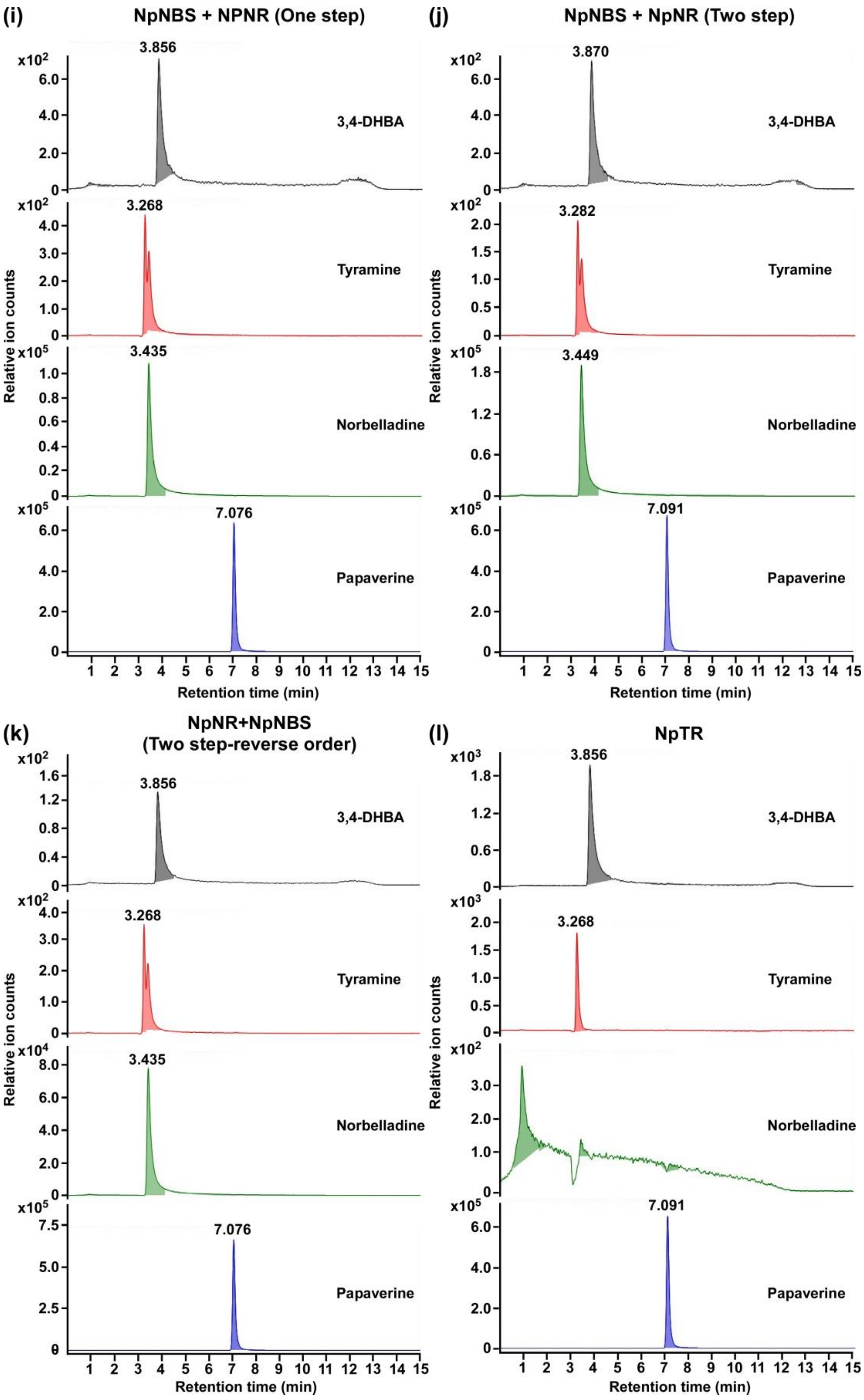
(i)
NpNBS + NPNR (One step)
(j)
NpNBS + NpNR (Two step)
(k)
NpNR+NpNBS
(Two step-reverse order)
(l)
NpTR
3,4-DHBA
Tyramine
Norbelladine
Papaverine
3.856
3.268
3.435
7.076
3.870
3.282
3.449
7.091
Relative ion counts
Retention time (min)

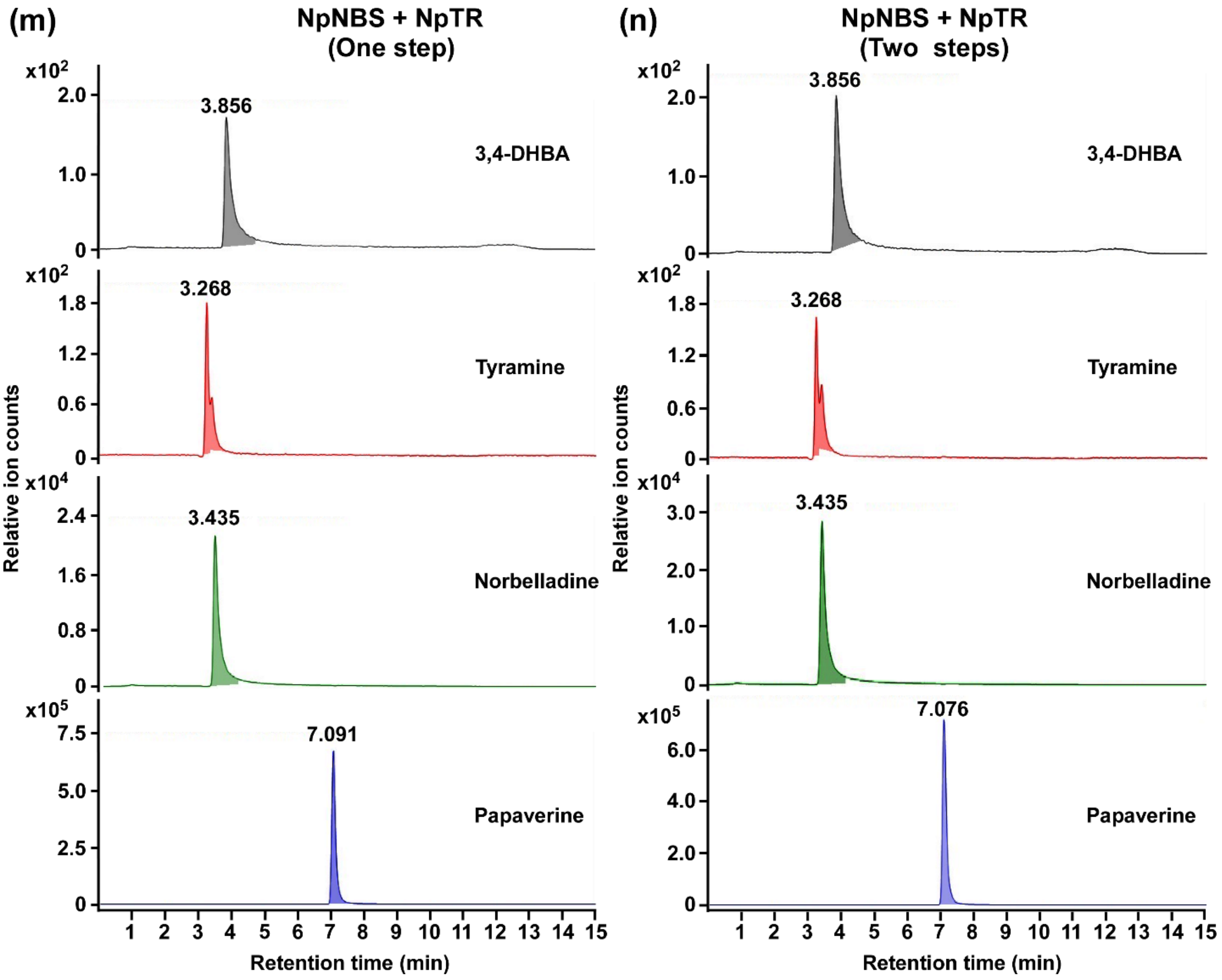
(m)
NpNBS + NpTR
(One step)
x10^2
2.0
1.0
0
3.856
3,4-DHBA
x10^2
1.8
1.2
0.6
0
3.268
Tyramine
x10^4
2.4
1.6
0.8
0
3.435
Norbelladine
x10^5
7.5
5.0
2.5
0
7.091
Papaverine
Relative ion counts
1 2 3 4 5 6 7 8 9 10 11 12 13 14 15
Retention time (min)
(n)
NpNBS + NpTR
(Two steps)
x10^2
2.0
1.0
0
3.856
3,4-DHBA
x10^2
1.8
1.2
0.6
0
3.268
Tyramine
x10^4
3.0
2.0
1.0
0
3.435
Norbelladine
x10^5
6.0
4.0
2.0
0
7.076
Papaverine
Relative ion counts
1 2 3 4 5 6 7 8 9 10 11 12 13 14 15
Retention time (min)

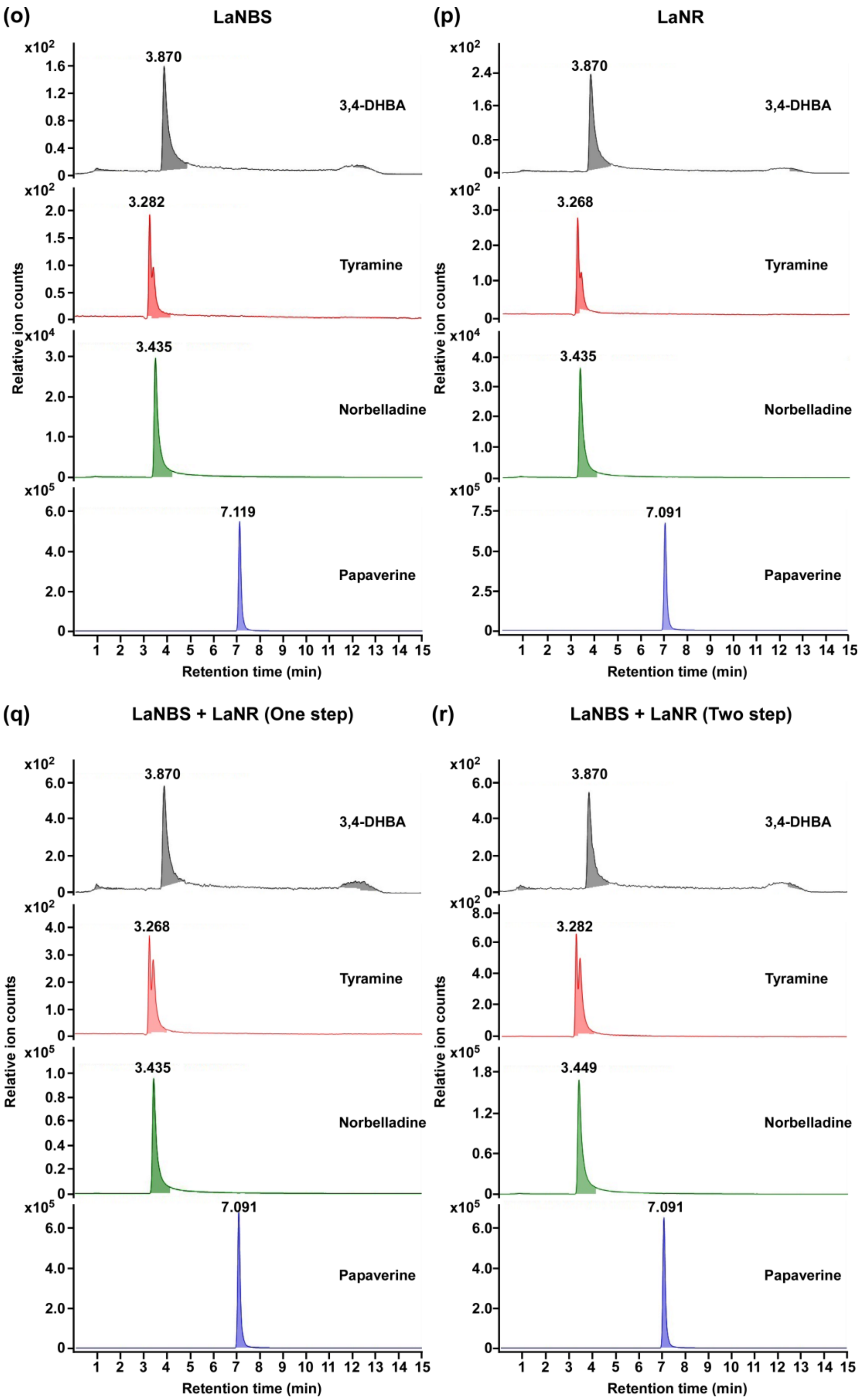

(o)
LaNBS
(p)
LaNR
3,4-DHBA
Tyramine
Norbelladine
Papaverine
3.870
3.282
3.435
7.119
3.268
7.091
Relative ion counts
Retention time (min)
(q)
LaNBS + LaNR (One step)
(r)
LaNBS + LaNR (Two step)
3.449

**Figure S9.** Extracted ion chromatograms of quantifier MRM transitions showing the detection of substrates, product, and internal standard of NBS and NR enzyme assays. Quantifier MRM transitions shown correspond to 260 → 138 m/z for norbelladine, 137 → 108 m/z for 3,4-DHBA, 138 → 77 m/z for tyramine and 340 → 202 m/z for papaverine respectively. The tested substrates used were tyramine (10 μM) and 3,4 dihydroxybenzaldehyde (300 μM). The extracted ion chromatogram corresponds to the mixed standards injected at 1 ppm (a), assays conducted without substrates (b), with the MBP tag alone (c), without enzyme (d), the complete assay performed with recombinant *Np*NBS (e), *Np*NBS assay with NADPH (f), the complete assay performed with recombinant *Np*NR (g), *Np*NR assay without NADPH (h), *Np*NBS and *Np*NR together one-step assay (i), *Np*NBS and *Np*NR together two-step assay (j), reverse order assay (k), *Np*TR assay (l), *Np*NBS and *Np*TR together one-step assay (m), *Np*NBS and *Np*TR together two-step assay (n), *La*NBS assay (o), *La*NR assay (p), *La*NBS and *La*NR together one-step assay (q), *La*NBS and *La*NR together two-step assay (r). Parent ion mass-to-charge (m/z) of 260 for norbelladine, m/z 137 for 3,4-DHBA, m/z 138 for tyramine, and m/z 340 for papaverine were subjected to collision-induced dissociation using multiple reaction monitoring analysis.

**Figure S10**

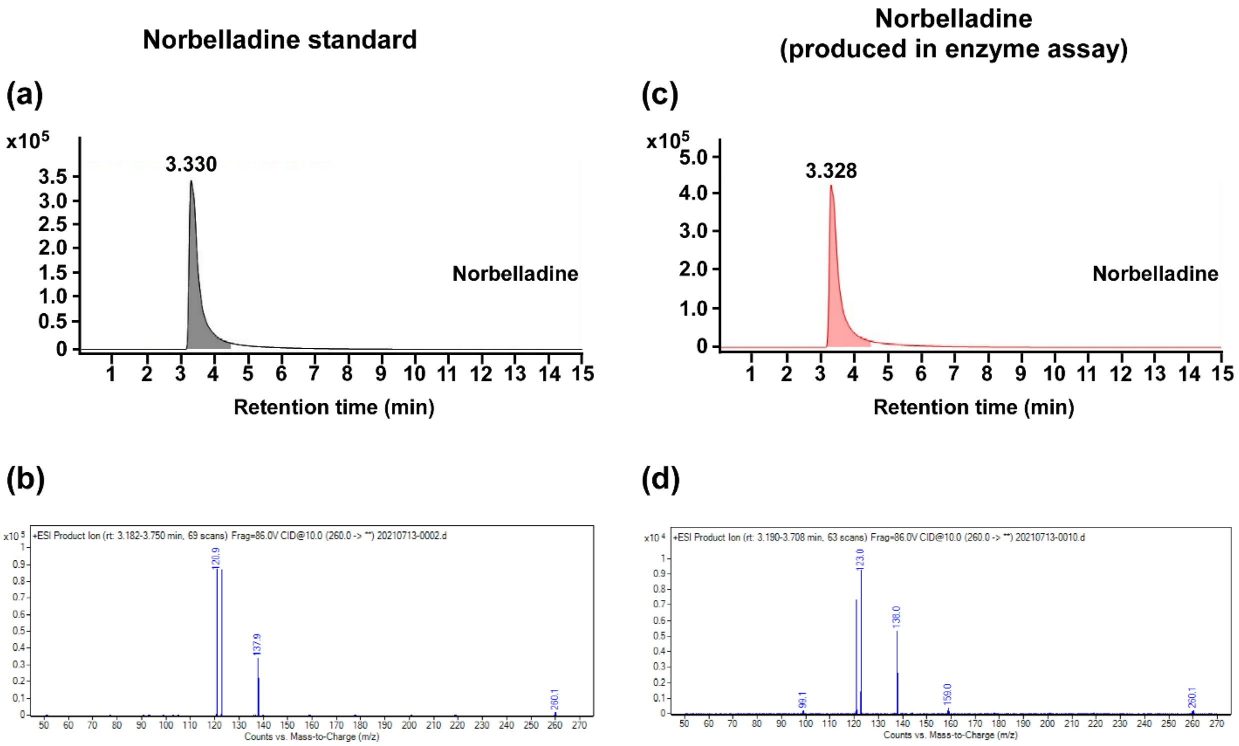


**Figure S10.** Comparison of the +ESI-MS/MS mass spectra of authentic norbelladine standard (a,b) and the product of the NBS and NR reaction (c,d). The extracted ion chromatogram corresponds to the norbelladine standard (a), or assay performed with recombinant *La*NBS (c). Parent ion mass-to-charge (*m/z*) of 260 for norbelladine was subjected to collision-induced dissociation analysis for identification and quantitation (b,d). Mass spectra of total ion chromatogram (TIC) obtained after the fragmentation using collision-induced dissociation (CID) of 10 electron Volts (eV) of the norbelladine standard (b) compared to the norbelladine produced by *La*NBS *in vitro* enzyme assay (d).

Figure S11

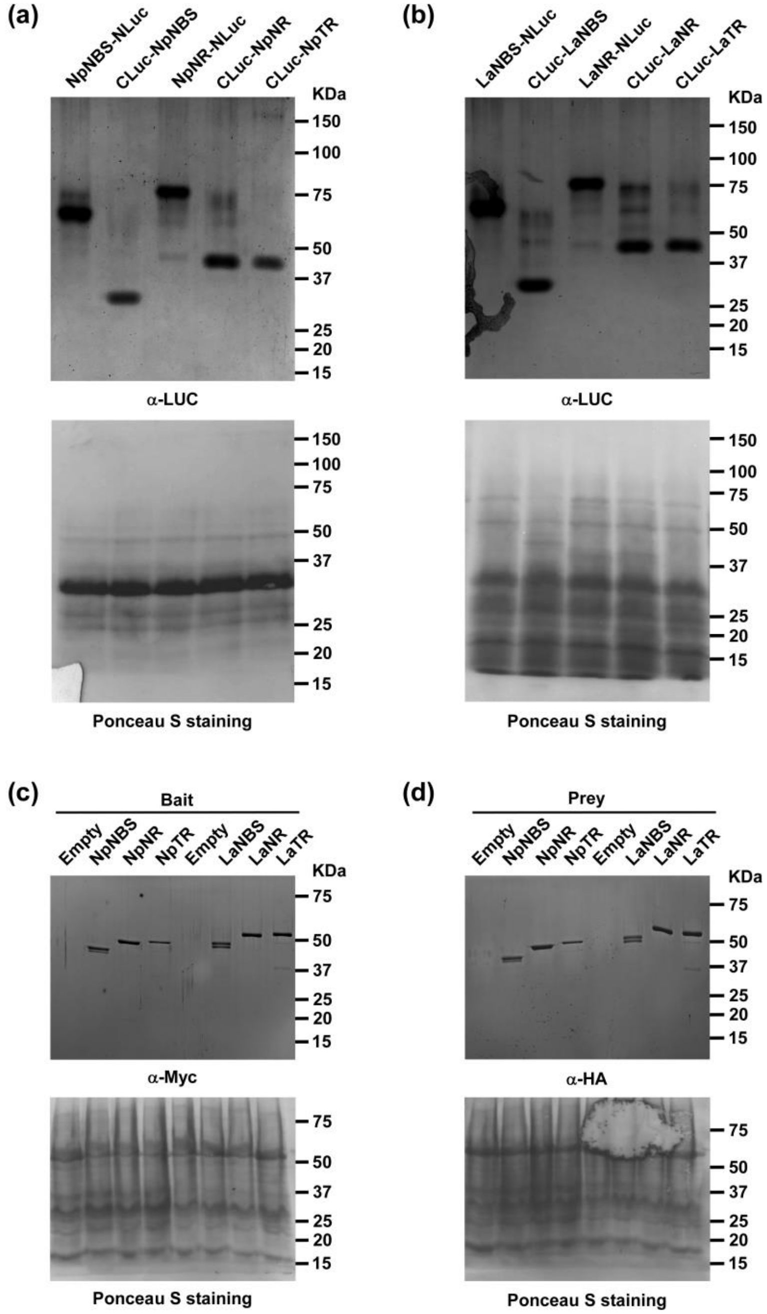


**Figure S11.** Protein expression of NBS and NR fusion *in planta* and in yeast. (a, b) Expression in *N. benthamiana* leaves of the indicated proteins fused to the N-terminal (NLuc) or C-terminal half (CLuc) of the luciferase protein detected by western blot analysis with anti-luciferase antibodies (α-LUC). Ponceau S staining of Rubisco is shown as a loading control. Numbers on the right refer to the location of standard protein molecular weight markers in kDa. (c, d) Expression in yeast of the indicated bait (fused to the GAL4 DNA-binding domain) and prey (fused to the GAL4 DNA-activation domain) proteins detected by western blot analysis using anti-MYC (α-Myc) and anti-HA antibodies (α-HA). Ponceau S staining of Rubisco is shown as a loading control. Numbers on the right refer to the location of standard protein molecular weight markers in kDa.

Figure S12

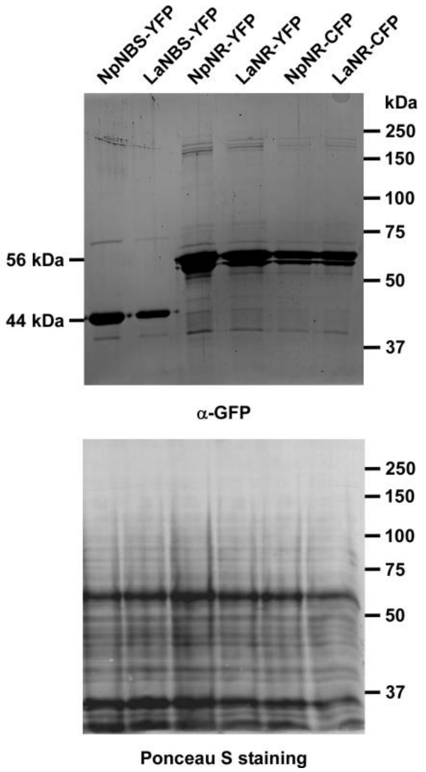


**Figure S12.** Protein expression of YFP- and CFP-fusion of NBS and NR *in planta*. Expression in *N. benthamiana* leaves of the indicated proteins fused to the C-terminal YFP or CFP detected by western blot analysis with anti-GFP antibodies (α-GFP). Ponceau S staining of Rubisco is shown as a loading control. Numbers on the right refer to the location of standard protein molecular weight markers in kDa.

Figure S13

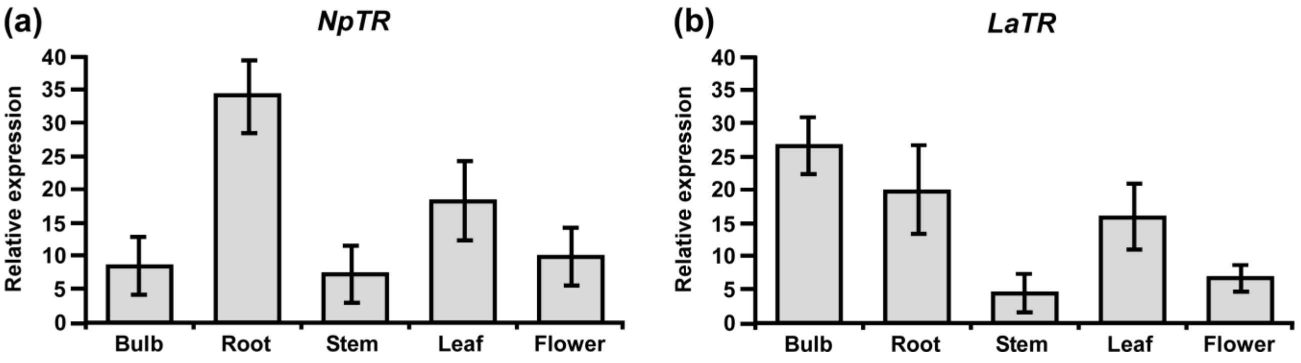


**Figure S13.** Relative expression of *TR* in different tissues of *N. papyraceus* and *L. aestivum* using reverse transcription quantitative PCR (RT-qPCR) analysis. Different plant tissues as indicated were harvested after flowering, *NpTR* (a), and *LaTR* (b) mRNA levels were measured by RT-qPCR analysis relative to expression in leaves. *NpHISTONE* and *LaGAPDH* were used as normalizer. Data are means ± *SE* of three biological repeats.

**Table S1.** Active site description and predicted interactions with ligands.

| | Active Site residues | Ligand | Score (kCal/mol) | Interactions Hydrophobic | H-bonds | Other |
|---|---|---|---|---|---|---|
| *Np*NBS | Tyr22, Leu27, Ala28, Ser31, Leu34, Leu35, Val38, Ile39, Val42, Leu55, Val57, Tyr59, Ile63, Pro64, Tyr68, Glu71, Phe73, Lys83, Thr85, Ile102, Phe104, Phe134, Asp138, Pro139, Phe140, Ile142 and Ile143 | Tyramine | -5,034 | Phe73, Thr85 | Ser31, Tyr59 | nd |
| | | 3,4-DHBAOP | -5,23 | Phe73,Thr85, Phe104, Ile143 | Glu71, Lys83 | nd |
| *La*NBS | Tyr22, Leu27, Ala28, Ser31, Leu34, Leu35, Val38, Ile39, Val42, Leu55,Val57, Tyr59, Ile63, Pro64, Gly65, Ile66, Lys67, Tyr68, His69, Glu71, Phe73, Lys83, Val85, Gly91, His92, Leu95, Tyr100, Ile102, Leu104, Phe134, Ala135, Thr136, Asp138, Pro139, Leu140, Ile142, and Ile143 | Tyramine | -5.10 | Leu31, Leu34 | Lys83 | nd |
| | | 3,4-DHBAOP | -4,72 | Val85, Leu204, Ile143, Leu34 | Lys83, Glu71, Tyr68 | nd |
| *Np*NR | Thr29, Gly30, Gly31, Thr32, Lys33, Gly34, Ile35, Gly36, Cys53, Ser54, Arg55, Asn56, Cys79, Asp80, Val81, Ser82, Asn108, Ala109, Gly110, Thr111, Ala112, Leu114, Lys115, Pro116, Thr131, Asn132, Ile158, Ser159, Thr160, Val161, Cys162, Leu167, Glu168, Gly169, His170, Tyr173, Lys177, Pro203, Gly204, Ile205, Thr206, Thr208, Val209, Gln210, Val211, Phe214, Leu215, Asp216, Asp217, Ala220, Lys223, Glu224, Arg227, Ile228, and Arg265 | NADPH | -11,51 | Ile35 | Lys33, Ile35, Gly36, Arg55, Cys79, Asn108, Gly110, Thr111, Lys177, Thr206, Thr208, Val209, Gln210 | π-cation Arg 55, Salt Bridge Arg 55 |
| | | Norcraugsodine | -6.36 | Leu114, Ile205, Phe214, Leu215 | Glu224, Arg265 | nd |
| *La*NR | Thr29 Gly30, Thr32, Lys33, Gly34, Ile35, Ser54, Arg55, Cys79, Asp80, Val81, Ser82, Asn107, Asn108, Ala109, Gly110, Thr111, Ala112, Ile113, Leu114, Lys115, Ile117, Ile120, Leu128, Thr131, Asn132, Phe133, Ser135, Ala136, Ile158, Ser159, Thr160, Val161, Cys162, Ile165, Leu167, Glu168, Gly169, His170, Ser171, Ile172, Tyr173, Ala174, Lys177, Met180, Pro203, Gly204, Ile205, Thr206, Thr208, Val209, Gln210, Val211, Phe214, Asp217, Asp219, Ala220, Lys223, Arg227, Ile228, Asp262, Arg265 and Gly266 | NADPH | -11,04 | Thr208, Gln210, Val211 | Lys33, Gly34, Ile35, Arg55, Asn56, Ser82, Asn108, Gly110, Asn132, Lys177, Thr208, Val209, Gln210, Val211 | Salt bridge : Lys33, Arg55 |
| | | Norcraugsodine | -6,15 | Ala112, Phe214 | Ala112, Gly160, Asp262 | nd |

| | | |
|---|---|---|
| *Np*TR | Thr28, Gly29, Gly30, Thr31, Arg32, Gly33, Ile34, Gly35, Cys52, Ser53, Arg54, Lys55, Cys78, Asp79, Leu80, Ser81, Asn106, Asn107, Val108, Gly109, Thr110, Asn111, Ile112, Arg113, Lys114, Pro115, Thr116, Tyr119, Phe126, Thr130, Asn131, Ile157, Ser158, Ser159, Val160, Ala161, Ala165, Val166, Asn167, Ser168, Gly169, Ser170, Tyr172, Lys176, Pro202, Trp203, Tyr204, Ile205, Thr207, Ser208, Leu209, Val210, Lys218, Ser219, Met220, Ser222, Ile223, and Arg226 | n.d. |
| *La*TR | Gly26, Gly27, Thr28, Arg29, Gly30, Ile31, Gly32, Ser50, Arg51, Lys52, Asp55, Asn103, Asn104, Val105, Gly106, Thr107, Asn108, Ile109, Arg110, Lys111, Asn128, Ile154, Ser155, Ser156, Val157, Ala158, Ala162, Ile163, Asn164, Ser165, Gly166, Tyr169, Lys173, Pro199, Trp200, Tyr201, Ile202, Thr204, Ser205, Leu206, Val207, Leu210, Leu211, Thr212, Asp213, Glu214, Lys215, Phe216, Val217, Glu218, Arg219, Val220, and Arg223 | n.d. |

**Table S2.** Oligonucleotides used in this study.

| Gene | Primer name | Sequence (5’→3’) |
|---|---|---|
| **Cloning into pMAL-c2x (protein expression in *E. coli*)** | | |
| *LaNBS* | LaNBS-FP-BamHI | AACGGGATCCATGAAGGGAAGTCTCTCCCATGAG |
| | LaNBS-RP-HindIII | ACGCAAGCTTCTACGCTACAATAGCTTTTTGCTCC |
| *NpNBS* | NpNBS-FP-BamHI | AACGGGATCCATGAAGGGAAGTCTGTCCCATGAG |
| | NpNBS-RP-HindIII | ACGCAAGCTTTCATGCTACAGTAGCTTTTTTCCGC |
| *LaNR* | LaNR-FP-BamHI | AACGGGATCCATGTTTACAGGAAGAGAAGAGGAATCAATGC |
| | LaNR-RP-HindIII | ACGCAAGCTTTCAACCGTTTATGCCCCGTCCTC |
| *NpNR* | NpNR-FP-BamHI | AACGGGATCCATGCTTAGAAGAAGAGAAGAGGAATCATTATC |
| | NpNR-RP-HindIII | ACGCAAGCTTTCAACCGTTTATGCCCCGTCCTCCGT |
| *LaTR* | LaTR-FP-BamHI | AACGGGATCCATGAAAGCAGCAGCAAAGAAAATCC |
| | LaTR-FP-SalI | ACGCGTCGACTTAATCGTGAGTTGGGTAGAAACCATTCAC |
| *NpTR* | NpTR-FP-BamHI | AACGGGATCCATGGGAGATAATGAGAGCAGCAGC |
| | NpTR-FP-SalI | ACGCGTCGACTTAATCATGAGTTGGGTAGAAACCATTCACGG |
| **Cloning into pBTEX-YFP (subcellular localization)** | | |
| *LaNBS* | LaNBS_FP_KpnI | AACGGGTACCATGAAGGGAAGTCTCTCCCATGAG |
| | LaNBS_RP_XbaI | ACGCTCTAGACGCTACAATAGCTTTTTGCTCC |
| *NpNBS* | NpNBS_FP_KpnI | AACGGGTACCATGAAGGGAAGTCTGTCCCATGAG |
| | NpNBS_RP_XbaI | ACGCTCTAGATGCTACAGTAGCTTTTTTCCGC |
| *LaNR* | LaNR_FP_KpnI | AACGGGTACCATGTTTACAGGAAGAGAAGAGGAATCAATGC |
| | LaNR_RP_XbaI | ACGCTCTAGAACCGTTTATGCCCCGTCCTCC |
| *NpNR* | NpNR_FP_KpnI | AACGGGTACCATGCTTAGAAGAAGAGAAGAGGAATCATTATC |
| | NpNR_RP_XbaI | ACGCTCTAGAACCGTTTATGCCCCGTCCTCCGT |

| | | |
|---|---|---|
| **Cloning into pCAMBIA1300:NLuc and pCAMBIA1300:CLuc vectors (SLCA assay for *in planta* protein-protein interaction)** | | |
| *LaNBS* | LaNBS_FP_nLUC_KpnI | AACGGGTACCATGAAGGGAAGTCTCTCCCATGAG |
| | LaNBS_RP_nLUC_SalI | ACGCGTCGACCGCTACAATAGCTTTTTGCTCC |
| | LaNBS_FP_cLUC_KpnI | AACGGGTACCATGAAGGGAAGTCTCTCCCATGAG |
| | LaNBS_RP_cLUC_SalI | ACGCGTCGACCTACGCTACAATAGCTTTTTGCTCC |
| *NpNBS* | NpNBS_FP_nLUC_KpnI | AACGGGTACCATGAAGGGAAGTCTGTCCCATGAG |
| | NpNBS_RP_nLUC_XhoI | ACGCCTCGAGTGCTACAGTAGCTTTTTTCCGC |
| | NpNBS_FP_cLUC_KpnI | AACGGGTACCATGAAGGGAAGTCTGTCCCATGAG |
| | NpNBS_RP_cLUC_BamHI | ACGCGGATCCTCATGCTACAGTAGCTTTTTTCCGC |
| *LaNR* | LaNR_FP_nLUC_KpnI | AACGGGTACCATGTTTACAGGAAGAGAAGAGGAATCAATGC |
| | LaNR_RP_nLUC_SalI | ACGCGTCGACACCGTTTATGCCCCGTCCTC |
| | LaNR_FP_cLUC_KpnI | AACGGGTACCATGTTTACAGGAAGAGAAGAGGAATCAATGC |
| | LaNR_RP_cLUC_SalI | ACGCGTCGACTCAACCGTTTATGCCCCGTCCTC |
| *NpNR* | NpNR_FP_nLUC_KpnI | AACGGGTACCATGCTTAGAAGAAGAGAAGAGGAATCATTATC |
| | NpNR_RP_nLUC_XhoI | ACGCCTCGAGACCGTTTATGCCCCGTCCTCCGT |
| | NpNR_FP_cLUC_KpnI | AACGGGTACCATGCTTAGAAGAAGAGAAGAGGAATCATTATC |
| | NpNR_RP_cLUC_BamHI | ACGCGGATCCTCAACCGTTTATGCCCCGTCCTCCGT |
| *LaTR* | LaTR_FP_cLUC_KpnI | AACGGGTACCATGAAAGCAGCAGCAAAGAAAATCC |
| | LaTR_RP_cLUC_SalI | ACGCGTCGACTTAATCGTGAGTTGGGTAGAAACCATTCAC |
| *NpTR* | NpTR_FP_cLUC_KpnI | AACGGGTACCATGGGAGATAATGAGAGCAGCAGC |
| | NpTR_RP_cLUC_SalI | ACGCGTCGACTTAATCATGAGTTGGGTAGAAACCATTCACGG |
| **Cloning into pGBKT7 (bait) and pGADT7 (prey) plasmids (yeast two-hybrid assay)** | | |
| *LaNBS* | LaNBS_SmaI | AACGCCCGGGATGAAGGGAAGTCTCTCCCATGAG |

| | | |
|---|---|---|
| | LaNBS_SalI | ACGCGTCGACCTACGCTACAATAGCTTTT TGCTCC |
| *LaNR* | LaNR_SmaI | AACGCCCGGGATGTTTACAGGAAGAGAA GAGGAATCAATGC |
| | LaNR_SalI | ACGCGTCGACTCAACCGTTTATGCCCCGT CCTC |
| *LaTR* | LaTR_SmaI | AACGCCCGGGATGAAAGCAGCAGCAAAG AAAATCC |
| | LaTR_SalI | ACGCGTCGACTTAATCGTGAGTTGGGTAG AAACCATTCAC |
| *NpNBS* | NpNBS_SmaI | AACGCCCGGGATGAAGGGAAGTCTGTCC CATGAG |
| | NpNBS_XhoI | ACGCCTCGAGTCATGCTACAGTAGCTTTT TTCCGC |
| *NpNR* | NpNR_SmaI | AACGCCCGGGATGCTTAGAAGAAGAGAA GAGGAATCATTATC |
| | NpNR_XhoI | ACGCCTCGAGTCAACCGTTTATGCCCCGT CCTCCGT |
| *NpTR* | NpTR_SmaI | AACGCCCGGGATGGGAGATAATGAGAGC AGCAGC |
| | NpTR_XhoI | ACGCCTCGAGTTAATCATGAGTTGGGTAG AAACCATTCACGG |
| **Primers used for RT-qPCR** | | |
| *LaNBS* | LaNBS_qPCR_FP | GCCCAGTGTCATCTCCAAGGT |
| | LaNBS_qPCR_RP | GAAAGCCAGATCCAAGTGCCC |
| *NpNBS* | NpNBS_qPCR_FP | CCCAGCGTCATCTCCAAGGT |
| | NPNBS_qPCR_RP | GTGCCCTCCCTCTACGATAAGTG |
| *LaNR* | LaNR_qPCR_FP | CAGTATCGTCCACATCTCCAC |
| | LaNR_qPCR_RP | ATGCCATCTCTAGCCCACTC |
| *NpNR* | LaNR_qPCR_FP | CATTTGAGCCAACTATCCCACCC |
| | LaNR_qPCR_RP | ATGCCGTCTCTAGCCCACTC |
| *LaTR* | LaTR_qPCR_FP | GTCAACTTGCACATCCTCTTCTG |
| | LaTR_qPCR_RP | ATGTTGTCCTTTGCCCATTCAC |
| *NpTR* | NpTR_qPCR_FP | GTCAACTTGCTCATCCTCTTCTG |
| | NpTR_qPCR_RP | GTTGTCCTTTGCCCATTCAC |
| *LaGAPDH* | LaGAPDH_qPCR_FP | GATCACTTTCCGTTCCAGGGTC |
| | LaGAPDH_qPCR_RP | CAGGGTGTAAGAGGGAGAAGG |
| *NpHISTONE* | NpHISTONE_qPCR_FP | GTCTGCCCCAACAACTGGAGG |
| | NpHISTONE_qPCR_RP | GCTTCCTAATCAGTAGCTCG |
